# Impact of symmetry in local learning rules on predictive neural representations and generalization in spatial navigation

**DOI:** 10.1101/2024.05.27.595705

**Authors:** Janis Keck, Caswell Barry, Christian F. Doeller, Jürgen Jost

## Abstract

In spatial cognition, the Successor Representation (SR) from reinforcement learning provides a compelling candidate of how predictive representations are used to encode space. In particular, hippocampal place cells are hypothesized to encode the SR. Here, we investigate how varying the temporal symmetry in learning rules influences those representations. To this end, we use a simple local learning rule which can be made insensitive to the temporal order. We analytically find that a symmetric learning rule results in a successor representation under a symmetrized version of the experienced transition structure. We then apply this rule to a two-layer neural network model loosely resembling hippocampal subfields CA3 - with a symmetric learning rule and recurrent weights - and CA1 - with an asymmetric learning rule and no recurrent weights. Here, when exposed repeatedly to a linear track, neurons in our model in CA3 show less shift of the centre of mass than those in CA1, in line with existing empirical findings. Investigating the functional benefits of such symmetry, we employ a simple reinforcement learning agent which may learn symmetric or classical successor representations. Here, we find that using a symmetric learning rule yields representations which afford better generalization, when the agent is probed to navigate to a new target without relearning the SR. This effect is reversed when the state space is not symmetric anymore. Thus, our results hint at a potential benefit of the inductive bias afforded by symmetric learning rules in areas employed in spatial navigation, where there naturally is a symmetry in the state space.

**Author summary:** The hippocampus is a brain region which plays a crucial role in spatial navigation for both animals and humans. Contemporarily, it’s thought to store predictive representations of the environment, functioning like maps that indicate the likelihood of visiting certain locations in the future. In our study, we used an artificial neural network model to learn these predictive representations by adjusting synaptic connections between neurons according to local learning rules. Unlike previous research, our model includes learning rules that are invariant to the temporal order of events, meaning they are symmetric with respect to the reversal of input timings. This approach produces predictive representations particularly useful for understanding spatial relationships, as navigating from one point to another is often equivalent to the reverse. Our model offers additional insights: it replicates observed properties of hippocampal cells and helps an artificial agent solve navigation tasks. The agent trained with our model not only learns to navigate but also generalizes better to new targets compared to traditional models. Our findings suggest that symmetric learning rules enhance the brain’s ability to create useful predictive maps for problems which are inherently symmetric, as is navigation.

## 1 Introduction

The hippocampus and its adjacent sub- and neocortical regions are widely believed to form both a crucial part in the acquisition and storage of memory, as well as the encoding of spatial and navigational variables in the form of spatially stable neural responses. (Scoville and Milner, 1957; O’Keefe and Nadel, 1978; Eichenbaum et al., 1999; Squire et al., 2004; Hafting et al., 2005).

It is an increasingly popular assumption that the representations that the brain generates in general, and in particular for space and memory, are not merely descriptive of the current state of the world, post-dictions of events or places just passed. Rather, it is believed that a predictive representation is learned, such that the objective is to infer future states of the world from one’s experience (Rao and Ballard, 1999; Friston, 2002; Stachenfeld et al., 2017; Russek et al., 2017; Behrens et al., 2018).

One framework that has extensively been used to describe this objective on the algorithmic level comes from reinforcement learning. The so called ‘successor representation’ (SR), or more broadly ‘successor features’ (SF) are a generalization of the well known value function, and are essentially a conditional expectation: Given the current state, they encode a (weighted) expectation of future values of a given function of the states of the world (Dayan, 1993; Barreto et al., 2017). If that function is simply an indicator of the states, then one obtains the SR, which thus roughly encodes how often states will be visited in the future.

Originally, the successor representation was proposed as an intermediate between ‘model-based’ and ‘model-free’ reinforcement learning (Momennejad et al., 2017; Gershman, 2018), allowing the storage of certain information about the transition structure under a given policy - hence, affording some generalization to different reward structures - while still being possible to learn with a efficient temporal difference (TD) learning algorithm (Dayan, 1993; Russek et al., 2017). Later work has also used the SR for different objectives such as option discovery (Machado et al., 2017b) and reward free exploration (Yu et al., 2023). More generally, maintaining a predictive representation might be a useful feature of intelligent agents (biological or artificial ones) that have to plan their behaviour (Carvalho et al., 2024).

In the hippocampal navigation literature, the successor representation view has been influential because apart from fitting well with the more general predictive brain hypothesis, it could explain non-trivial effects of place cells that had been previously observed, for example the skewing of place fields in direction of travel or the non-extension of place-fields through obstacles in the environment (Stachenfeld et al., 2017). Furthermore, successor representation theory yielded an algorithmic explanation for grid cells as an eigendecomposition of place-cell structure, which could also be connected to efficient neurally plausible navigation (Dordek et al., 2016; Corneil and Gerstner, 2015; De Cothi and Barry, 2020; De Cothi et al., 2022).

Despite the success of SR theory explaining neural data on an algorithmic level, there has been considerably less work dedicated to providing a mechanism through which the SR should be learned using biologically plausible learning rules (Vértes and Sahani, 2019). Recently, this question has been tackled by the community: Two papers (George et al., 2023b; Bono et al., 2023) used feed-forward networks to learn synaptic weights that compute successor features from their inputs. (George et al., 2023b) focuses on the predictiveness afforded by the theta cycle together with a compartment-neuron learning rule, while (Bono et al., 2023) uses spiking neural networks and a spike-time-dependent plasticity (STDP) rule. On the other hand, (Fang et al., 2023) used a recurrent neural network, to learn successor features directly in the activities of the recurrently connected neurons.

Anatomically, the latter approach can be linked to plasticity occurring at the recurrent synapses of CA3, while the former approach maps on the feedforward synapses to CA1 (Knierim, 2015). Both areas are known to show a considerable proportion of place cells (Leutgeb et al., 2004), hence both are indeed candidate regions to encode successor representations. However, it has been suggested that different learning rules might be in place at the respective synapses: the Schaffer collateral-synapse to CA1 pyramidal cells is classically believed to obey the rules of STDP (Wittenberg and Wang, 2006; Markram et al., 2011), which in its stereotypical form requires presynaptic increased activity to precede postsynaptic increased activity for an increase in strength of synaptic connection (Markram et al., 1997; Bi and Poo, 1998). On the other hand, recent work has identified a regime in which recurrent CA3 synapses get strengthened if pre- and postsynaptic increased activity are close in time, regardless of the temporal order - and computationally linked a symmetric learning rule to benefit in memory storage of a recurrent network (Mishra et al., 2016).

Here, we want to investigate the effect of such symmetric learning rules on the construction of predictive representations. That is, we aim to understand whether using a learning rule insensitive to the temporal order of the inputs learns different successor representations - and which (dis-)advantages it yields. To this end, we first construct a model which has both a recurrent and a feedforward component, reminiscent of the architecture of Hippocampus, and study the successor features that are learned using a local learning rule. Thereby we extend the earlier work which focused on learning in a single layer to learning at multiple levels. This extension is rather straightforward and results in both layers learning successor features based on their respective inputs.

We then find that by changing the learning rule, the representations also undergo a similar modification: In the symmetric setting, instead of encoding future expectations under the current true policy of the agent, successor features under a symmetrized version of the transition probabilities are learned, while an asymmetric rule learns the ‘true’ successor features. We then contrast the utility of the respective representations in a reinforcement learning setting. There, we find that a symmetric learning rule yields benefits for generalization in navigational tasks, where the symmetry of the state space can be exploited, while an asymmetric learning rule is more advantageous for generalization in asymmetric state spaces. We conclude that implementing both an asymmetric and a symmetric learning rule might yield complementary representations.

## 2 Results

### 2.1 Successor Representations

The successor representation and the more general successor features describe future expectations of a quantity, conditional on the current state of the world. They are most easily defined in the following setting: Assume the environment of an agent/animal consists of a set of states 𝒮. The states of the world are changing according to a time homogeneous Markov chain, denoted *S*_*t*_ ∈ 𝒮, with transition probabilities encoded in the matrix *P* such that *P*_*s,s*_′ := *p*(*s*′| *s*), the probability transitioning from state *s* to state *s*′ in one timestep. Then for any feature/observation function

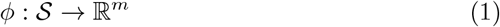

one can define an expectation of weighted, cumulative future values of that function, given the current state:

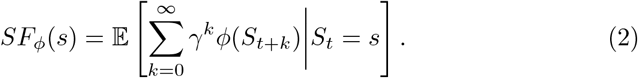

The weighting factor *γ* ∈ [0, 1) puts relatively more importance on proximal times. In the case that *ϕ* is an injective function, that is for every state of the world there is an unique observation value, it makes sense to define the ‘successor representation’ (SR)

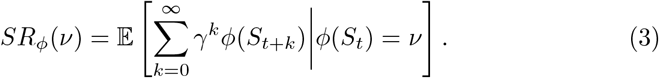

We use this terminology here in a little more generality than is usual, but it can be seen that one important special case leads to what is usually called SR: Let *ϕ*^*e*^(*s*) := *e*_*s*_, where *e*_*s*_ is the unit vector with a 1 at the entry corresponding to state *s*. That is, *ϕ*^*e*^ assigns to each state its indicator vector. Then one obtains

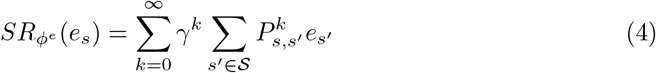

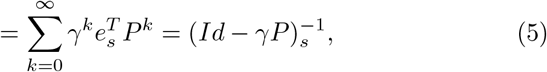

where the last equality is the well known identity for the Neumann series. The matrix (*Id* −*γP*)^−1^ is widely referred to as the SR, so our definition encompasses this special case. One can see from these equations that the SR of indicator vectors thus gives a weighted sum of expected future visitation probabilities of states, and it is this expression that has originally been used to model the predictive representations that hippocampus ought to encode (Stachenfeld et al., 2017; Russek et al., 2017).

### 2.2 Learning Successor representations with local learning rules

We construct a simple model of two neuron population activities *p*_*t*_ which have dynamics of the form

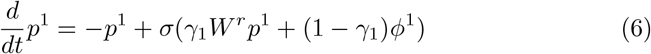

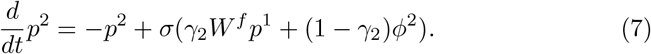

Here, *W*^*r*^ is a recurrent connectivity matrix that feeds the activity of the first population back into itself, while *W*^*f*^ is feedforward matrix, which encodes how activity of the first population is fed into the second. This architecture of one population of highly recurrently connected neurons feeding into a second one with little recurrent connectivity is reminiscent of hippocampal subfields CA3 and CA1 respectively (Knierim, 2015). The two populations obtain additional external inputs *ϕ*^1^, *ϕ*^2^, which might represent the input to CA3 via mossy fibers, or directly through the perforant path, and the input to CA1 from EC through the latter, respectively. In the simplest case, these inputs are just indicator-functions for particular states, that is *ϕ*_*i*_(*s*) = *δ*(*s* = *s*_*i*_). A more realistic shape, which we employ in our experiments in section 2.5, might be given by Gaussian inputs of the form 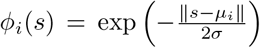. The *γ*_*i*_ ∈ [0, 1] are scalar valued global gain factors which control the relative strengths of inputs to the populations. Note that we intentionally used the same symbol *γ* as for the timescale factors in the successor representation, as those will turn out to be equivalent. For analysis, we assume that the activation function *σ* is the identity (i.e. the system is linear), and that the population vectors take the equilibrium values of the above dynamics, that is

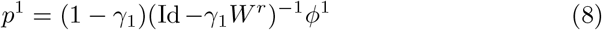

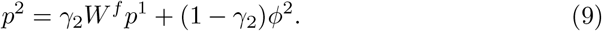

We then define a learning rule for the synaptic weights, using these equilibrium values. Hence, we implicitly assume that neural dynamics happen on a timescale *τ*_*p*_ which is way quicker than that of learning, *τ*_*W*_. Taking the equilibrium values is the limit case which simplifies analysis, but in practice one can also take *τ*_*p*_ *<< τ*_*W*_ and simply update activities and weights concurrently. We also assume that our weight matrices are initialized in such a way that these are stable equilibria of the dynamics, which for example will be the case if all weights are initialized to sufficiently small non-negative values.

The learning rule we use is a slight modification of the learning rule used in Fang et al. (2023). In particular, we use the same general learning rule for both recurrent and feedforward weights, only varying certain parameters. Let *p*^*post,i*^, *p*^*pre,i*^ be the activity of the *i*-th post/pre-synaptic neuron respectively. We then update the weight from the *j−* th presynaptic neuron to the *i*-th post-synaptic neuron via

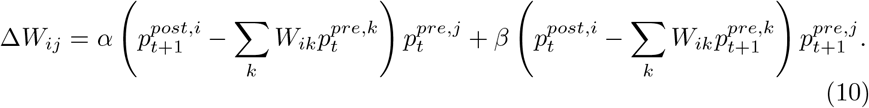

The update rule contains terms of the form *p*^*post,i*^*p*^*pre,j*^, which are simply Hebbian terms. The other summands perform a subtractive normalization: they subtract the total overall input to a post-synaptic neuron, such that only activity exceeding this input will actually be considered positive. This term has been interpreted as a decorrelative term by (Fang et al., 2023) - in total, the learning rule can be understand as a predictive coding rule approximating a conditional expectation operation, as we explain in appendix A. In matrix-vector notation the update rule reads

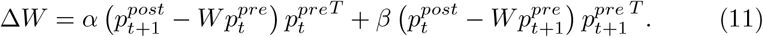

The parameters *α, β* ∈ ℝ control the sensitivity of this update to the order of the activity, put differently, the temporal symmetry of the rule: If we write our synaptic weight change as 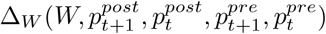, then for parameters *α* = *β* we obtain a learning rule that is invariant under a reversal of time, that is

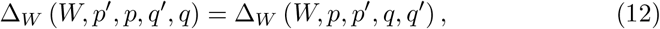

while for the original learning rule with *α* = 1, −*β* = 0, no such relation holds - see also Figure 1. For *α* = *β* we would obtain a rule that is antisymmetric in this sense - it turns out however that this would yield unstable dynamics.

**Figure 1:**
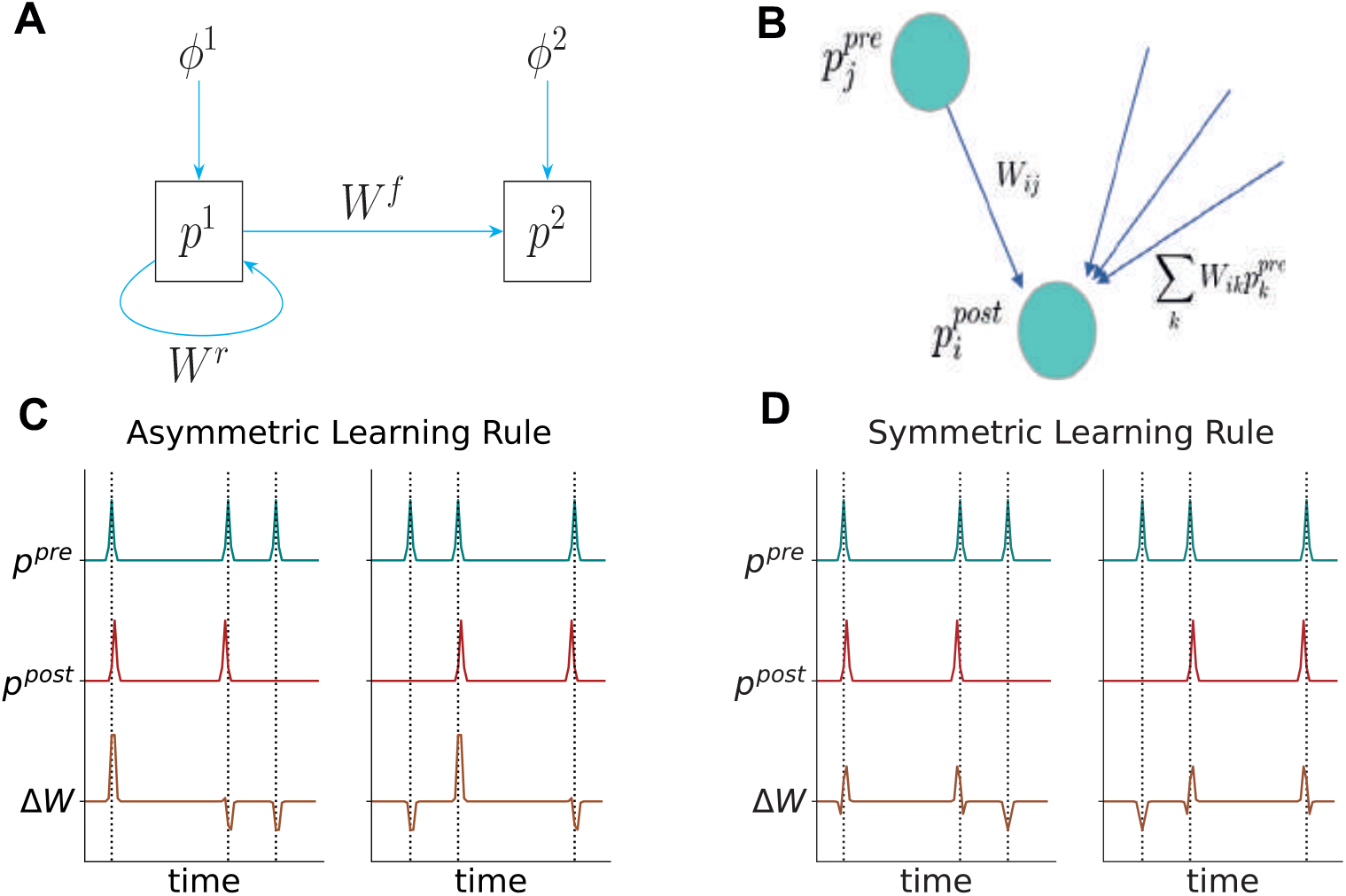
Model and learning rule. **(A)** Cartoon depiction of the model we are using in the main text. A recurrently connected population of neurons *p*_1_, putatively *CA*3, receives external input *ϕ*_1_, putatively from dentate gyrus or entorhinal cortex(*EC*). It projects to another population *p*_2_, which receives input *ϕ*_2_. The latter could be *CA*1 receiving input again from *EC*. Note that there are no recurrent connections in the second layer and no backwards connections. **(B)** Quantities relevant for the update of synapse *W*_*ij*_: pre- and postsynaptic activities, as well as the sum of the total input to the postsynaptic neuron through the synapses *W*. **(C), (D)** Invariance of learning rules with respect to temporal order. We plot synaptic weight change of a single synapse in a setup with a single pre- and postsynaptic neuron, respectively. The right column has the same pre- and postsynaptic activities as the left column, only in reverse order. In **(C)**, the learning rule with parameters *α* = 1, *β* = 0 is used, while in **(D)** 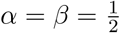. Only in the latter the synaptic weight changes are preserved (in reverse order), while in **(C)**, postsynaptic activity before presynaptic activity leads to a net weight decrease. Note that in this illustrative example *W* is fixed, in reality, network dynamics and weights would influence each other and lead to more complex changes.

### 2.3 Network learns successor representation and successor features

Before exhibiting the representations that are learned under the modified learning rule, it might be helpful understanding which representations a two layer-network as the above learns with the simplest choice of parameters (*α* = 1, *β* = 0). Let us assume the network has been exposed extensively to features *ϕ*^1^, *ϕ*^2^ under the same random walk with transition probabilities *P*, such that the synaptic weights could converge. In practice this means simulating discretized dynamics of Equation 6 together with the learning rule of Equation 10, which then results in updates of the form (*ε*_*p*_ *>> ε*_*W*_):

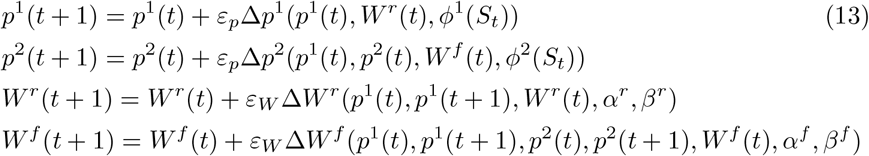

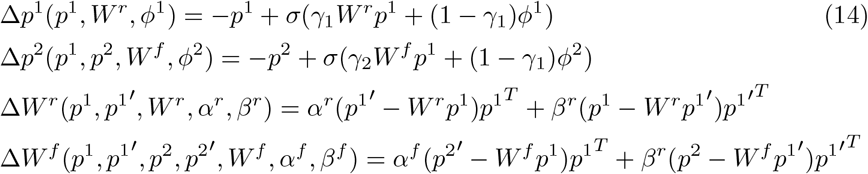

Then, as we show in appendix A and appendix D, the synaptic weights converge in such a way that the network computes successor features. Indeed, the equilibrium is best explained by stating what the population activities encode once the weights have converged. We find that the activities in the network, after extensive exposure to features *ϕ*^1^, *ϕ*^2^, when presented with new inputs 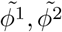, compute successor representations/features: They take the equilibrium values

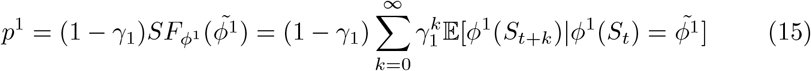

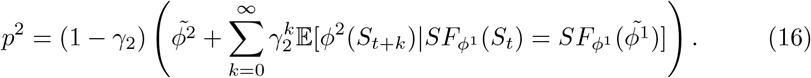

In words, this means that the first, recurrent layer computes the weighted cumulative sum of the predicted values of the feature *ϕ*^1^ it was trained on, but given the possibly new feature 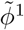 - one could see this a form of pattern completion, but with a predictive component. Similarly, the second layer computes predictions of *ϕ*^2^, only that these predictions themselves depend on those predictive representations passed to it from the first layer. In particular, when the inputs are the same as the model was trained on, and the maps *ϕ*_*i*_ are injective (i.e., sufficiently rich features exist for the environment), then these equations simplify to

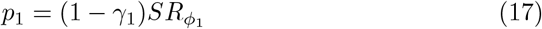

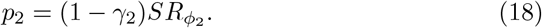

That is, in this case the two layers simply learn to encode the successor representations of their respective inputs.

#### 2.4 Influence on the representations by choice of parameters

We now proceed to ask the question “which representations would be learned depending on the choice of the parameters *α, β*?”. It turns out that the resulting representations are still successor representations, albeit corresponding to transition probabilities that are not necessarily faithful to those of the environmental dynamics anymore. To be precise, we define a weighted sum of transition matrices

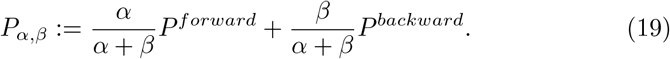

Here, *P*^*forward*^ contains the transition probabilities of the actually observed process (that is, ‘forward’ in time), while *P*^*backward*^ contains the transition probabilities of the reverse process, i.e. *p*(*s*_*t*_ = *s*| *s*_*t*+1_ = *s*′) - see also Equation 39 We then show in appendix D that under suitable conditions, with the learning rule defined above, the model is able to learn the successor representations under *P*_*α,β*_. This entails that the neuronal population activities at convergence yield

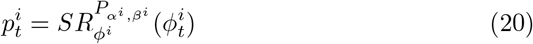

with *α*^*i*^, *β*^*i*^ the respective parameters used in the learning rule. In terms of the successor representation matrix, this simply means

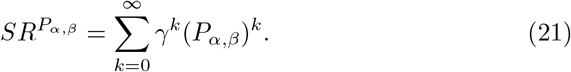

That is, in particular, under the regime *α* = 1, *β* = 0, this corresponds to the ‘true’ successor representation. For *β* = 1, *α* = 0 one obtains the ‘predecessor representation’ (Yu et al., 2023). Under a symmetric regime, the transition probabilities forward and backward in time are averaged over, that is one obtains

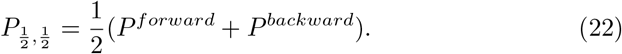

In fact, in this case the transition probabilities are *reversible* in time - this is not surprising, as the learning rule was defined to be invariant under a time reversal and hence should only extract aspects of the dynamics which are reversible. We note here that such reversible dynamics have been typically assumed in the theory of SR when construing grid cells as efficient representations of the geometry of an environment through the eigenvectors of the SR - see Appendix C.

##### 2.4.1 Activities in the model converge to theoretically obtained limits

Having theoreticallly obtained the limits of the weights and the corresponding activities, we next verified these limits empirically in simulations. In these simulations, we consider an environment with a discrete number of states *s*∈𝒮 and inputs *ϕ*^*i*^ which are functions of these states. In the simplest case, *ϕ*(*s*) = *e*_*s*_, we obtain the classical successor representation. In particular, for a symmetric learning rule we obtain a symmetrized successor representation - which shows less dependence on the policy. For example, on a circular track, the representation becomes indifferent to whether the agent is performing a clock-wise or a anti-clockwise walk (Figure 2). One might argue that a reversible representation encodes more of the geometry of the underlying state space and less of the actual dynamics (although there is still an indirect influence of the dynamics through the stationary distribution). Indeed, we show in appendix G that the symmetrized transition probabilities are always closer to a uniform policy than the unsymmetrized ones.

**Figure 2:**
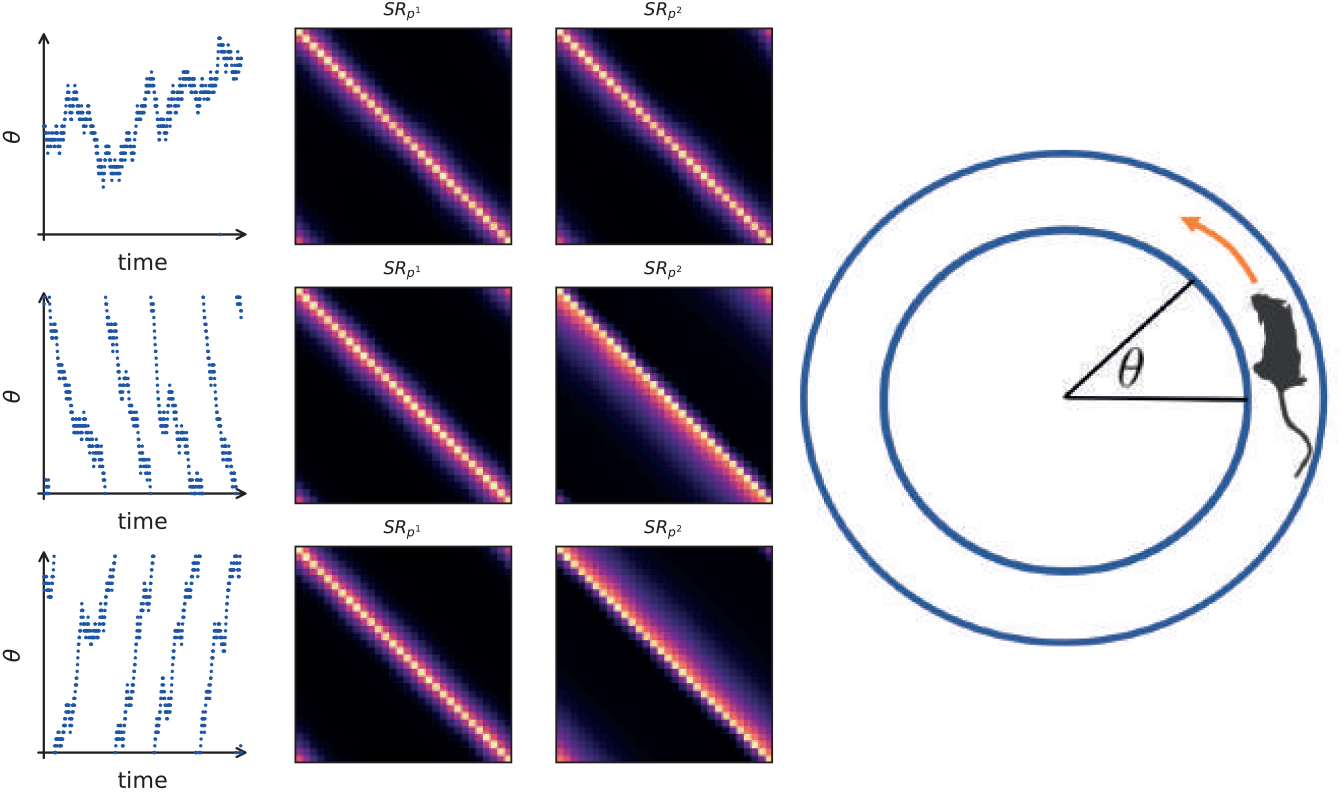
Successor Representations learned in circular random walks. We construct a circular state space with possible actions stay, move clockwise and move anti-clockwise. We simulate three random walks, one where the actions are selected uniformly (first row), one where clockwise actions are preferably selected (second row) and one where anti-clockwise actions are preferably selected (third row). The first column shows an example trajectory of the respective walk. The second and third column show the successor representations learned by the first and second layer of our model, using a symmetric (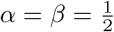 and an asymmetric (*α* = 1, *β* = 0) learning rule, respectively. Note how the successor representation learned with a symmetric rule does not distinguish between the policies. Here, the inputs to the cells are one-hot vectors encoding the respective states and the plotted successor representations are obtained by taking the average population activity in the respective states.

We also verified our theoretical results in more complex scenarios: the convergence prevails also when the features are random inputs instead of one-hot vectors, and also when the random walk is arbitrary instead of circular - see Figure 3. Additionally, we investigated the stability under the choice of parameters *α, β*. Here we found that it seems that the model is only stable when the positive weight is bigger (in absolute value) than the negative weight, with no convergence at the boundary case of *α* = − *β*. The latter is in line with the theoretical results - note that Equation 19 is undefined in this case.

**Figure 3:**
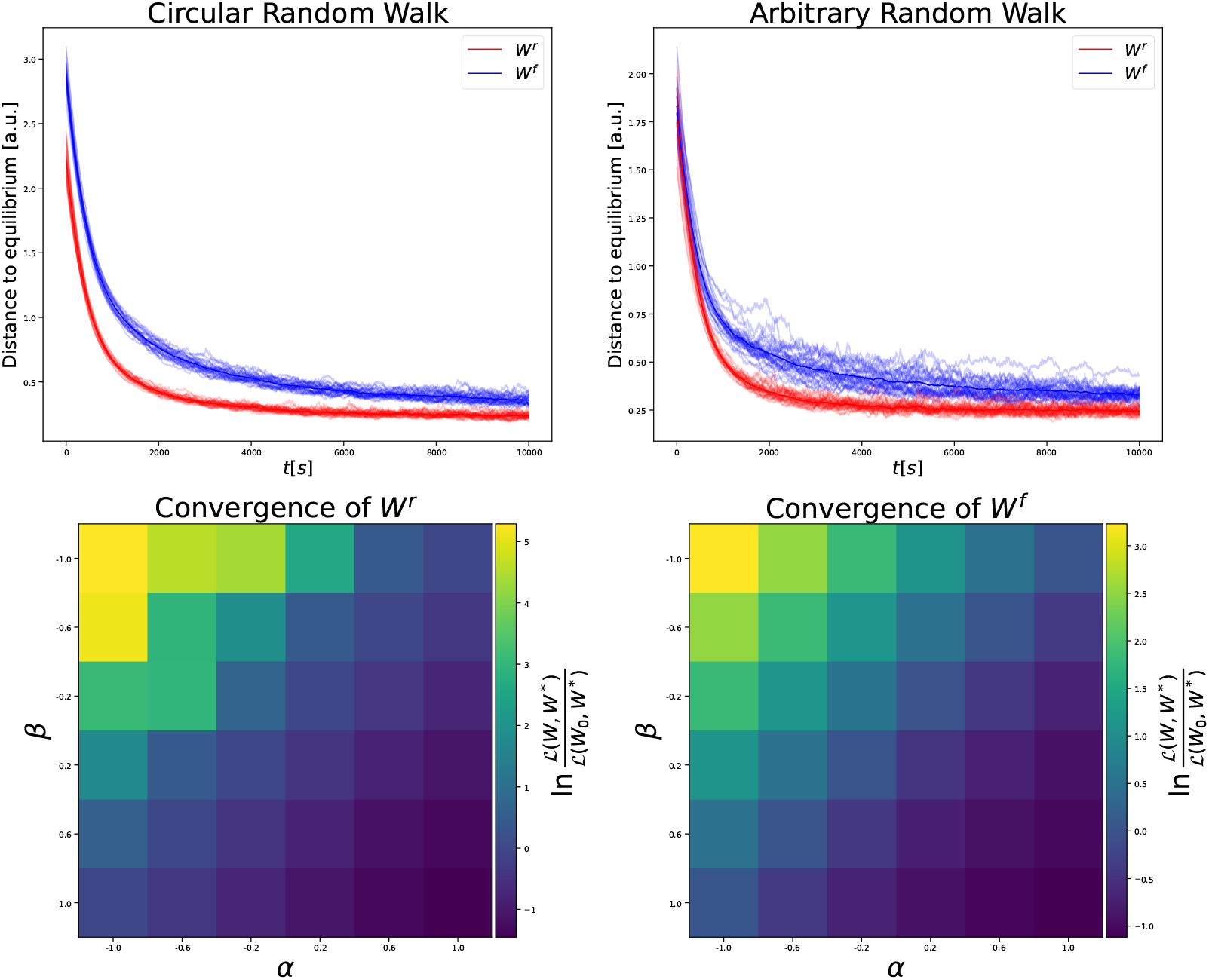
Successor representations are learned for a variety of inputs, dynamics, and parameters. **Top:** Convergence of recurrent (red) and feedforward (blue) matrices to their theoretical limit with random features in circular (**left**) and arbitrary (**right**) random walks. **Bottom:** Convergence of recurrent weight (**left**) and feedforward weight (**right**) for different parameters *α, β*. The other set of parameters is fixed to (1, 0) and 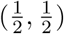 in these experiments, respectively. In graphs, we measure convergence by the loss term ℒ as explained in Methods section. In the bottom row, we compute the fraction of the loss at the final step over the initial loss and display the result in a logscale. Thus, negative values indicate converging towards the target. Note that the values on the antidiagonal are approximately 0.

### 2.5 Place fields under symmetrized rule show less shift

Although both areas CA3 and CA1 show place cells, these cells exhibit different properties and dynamics (Lee et al., 2004a,b). It is well known, that on a linear or circular track place fields shift backwards opposite the direction of travel in both regions (Mehta et al., 1997; Roth et al., 2012). However, when directly comparing cell recordings from both, it has been observed for example in Dong et al. (2021) that the shift in CA3 is in general less pronounced, that is, the center of mass of these cells is more stable than in their counterparts in CA1. We hypothesized that a difference in learning rules could explain this effect. Indeed, it is not hard to see theoretically why this should be the case: Through learning, the features *ϕ*^*i*^(*s*) get replaced by their successor features 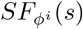. If there is a preferred direction of travel, then states preceding those where the feature puts a lot of mass will also have more mass, since they are predictive of the former states. If there is an asymmetry in the policy, the same will not hold true for succeeding states, hence one observes a shift towards the predecessors. If now however one has symmetric transition probabilities, then there is no directionality, hence this shift would not occur. Indeed, we provide a simple proof of this in subsection 5.5

We confirmed this intuition, running our two-layer model in a simple linear track where the agent repeatedly moves from the left side to the right. Indeed, we find a tendency to shift in the CA1 cells, which isn’t as pronounced in the CA3 population. Qualitatively, our results match those obtained in Dong et al. (2021). Importantly, these results only hold when using the symmetrized version of the learning rule for CA3, while the asymmetric variant yields almost no distinction.

### 2.6 Generalization and learning with (a)-symmetric rules

Having derived the different kinds of successor representations that symmetric and asymmetric learning rules encode, we next sought to understand what the functional benefits of these representations might be. In particular, we hypothesized that a symmetric learning rule for successor representations might be a relatively simple inductive bias that would favour learning such representations that are invariant under time reversal. This could be useful in such environments where there is a symmetry in transition structure. A particularly simple example of this setting but still likely for biological agents to encounter is when the transition structure of the environment is deterministic and transitions in both directions between states are possible. In this case, the state space becomes a metric space and the metric a particular invariant under the symmetry - that is *d*(*s, s*′) = *d*(*s*′, *s*). This then hints at a possible benefit of using a learning rule biased towards such symmetry: In a metric space, an optimal policy for navigating towards any target depends only on the metric - in fact one may give a closed form expression as we show in appendix G.1. Hence, the more the representations that are learned in one particular tasks encode something akin to the distance on the underlying space, the more useful these representations should be for generalizing to new such tasks. In other words, one would want to bias the representations to encode more of the geometry of the space and less of the dynamics of the particular tasks.

In the light of this hypothesis, we trained a reinforcement learning agent, equipped with a temporal difference (TD) learning rule that for a fixed policy would converge to the same weighted representation under *P*_*α,β*_, and investigated performance in simple navigation tasks. Note that since we now want to understand the benefits on the computational level irrespective of the biological implementation, we use a classical RL model and not the neural network model from the preceding sections - however, all experiments could also be conducted using such a model. The agent we use encounters transitions of states and updates an internal matrix *M* which serves as an estimate of the successor representation. When transitioning from state *s* → *s*′, the update equations are given by

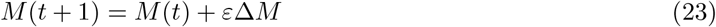

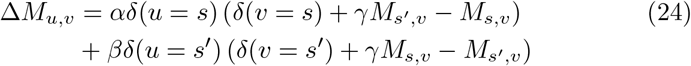

where *ε* is a fixed learning rate. Note that for *α* = 1, *β* = 0 this yields the standard TD learning rule for the successor representation (Gershman, 2018). Furthermore, the agent learns a reward vector *R*, and computes the value function of states via *V*_*R*_(*s*) = *MR*(*s*). Together with a local transition model *p*(*s*′| *s, a*) (which we assume as given), the agent can then define *Q* values of state action pairs *Q*(*s, a*) and take the next action based on these *Q* − values (Sutton and Barto, 2018; Dayan, 1993).

The task for the agent is split in two parts: In the first part, in each epoch, the agent is initialized in a random location and has to navigate to a fixed goal *s*_*target*_, where a unit reward is received -i.e., 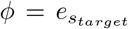. This goal does not change over epochs. After a fixed number of epochs, the goal is changed to a new location 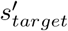, randomly drawn from all other locations. Importantly, the agent is then only allowed to relearn the reward vector, not the successor representation matrix.

We find that both the classical SR, corresponding to parameters *α* = 1, *β* = 0, as well as the symmetrized version 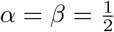 are able to learn the navigation tasks with similar mean learning curves. Although it might be counterintuitive at first that symmetrization yields a policy which still navigates to the correct target, we actually prove in appendix G.1 that indeed an optimal policy is stable under such symmetrization - which shows at least that once such a policy is learned, it can be maintained. However, this result only shows stability once an optimal policy is reached, and indeed one may observe in Figure 6 that the symmetric agent generally shows a higher variation. Furthermore, we find that the symmetric agent seems to be more sensitive to the choice of hyperparameters: Indeed, we find that for smaller learning rates, there is a steeper degradation of performance caused by high variability in the results for the symmetric agent. However, for higher learning rates, both show a similar performance.

Importantly, we then find empirically that on the new targets, the symmetric learning rule outperforms the classical one on average, while both show higher variation in these tests (Figure 6). Thus, one may argue that the successor representation in a symmetric learning regime affords better generalization - at least in a navigational setting. This is not merely an effect of the trajectories that are sampled with the different learning rules - that is, in particular it cannot be attributed to the higher variation during training on the first target: We repeated the above experiment while learning the successor representations based on the transitions obtained from the classical agent alone - the results remain unchanged, suggesting that the symmetric rule yields representations more apt to generalization without the need of a different sampling regime (Figure H.5). Similarly, the results also hold when controlling the norm of the updates, such that both the asymmetric and symmetric update make an equally big update step at each point. In other words, the agent can concentrate on solving the current task and still gets afforded a map of the environment which is less influenced by the current policy. Nevertheless, when learning to navigate to the first target, the symmetric agent still learns a policy that on average has more entropy than the one learned by the asymmetric agent. This increase in entropy then yields a the generalization advantage: as soon as the new target is introduced, the effect is reversed and the asymmetric agent has the more entropic policy (Figure 5).

Indeed, one can show that the transition probabilities encoded in 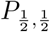 will be closer (than the observed transition probabilities) to those that correspond to a uniform policy, choosing every transition with equal probability - see appendix G - in particular meaning they have higher entropy. The successor representation of the uniform policy in turn is closely related to the shortest-path distance (Zhang et al., 2021). It can thus be used to generalize to *any* navigational target, while the successor representation under other policies will not neccesarily have this property. Together with our previous considerations on the symmetry of the state space, this led us to hypothesize that the generalization effect of the symmetric learning rule should vanish as soon as there is no such symmetry in the state space any more. We thus repeated the above experiment on a state space that corresponds to a directed graph, where the number of transitions needed to go from *s* to *s*′ is not necessarily equal to those needed to travel from *s*′ to *s*. Indeed, we find that in such a setting the effect is reversed: there, the classical learning rule leads to better generalization (Figure 7).

### 2.7 Variation in Symmetry

So far the temporal difference learning rules we used had either perfect symmetry or no symmetry at all. In biological agents, such perfect symmetry might rarely be given, rather, one might expect that the temporal sensitivity profile of learning rules could vary both in space and time, meaning that cells exhibit different learning rules at different moments and furthermore two cells might vary in how they learn. We wondered how robust our findings are to such variations. Thus, we investigated the generalization performance of our successor representation agents under variations that might correspond to imperfect symmetry. First, we investigated how well agents would generalize that have a learning rule in between the symmetric and the classical ones we have discussed so far. To do so, we chose parameters 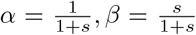, where *s* is a parameter that we varied. Note that for *s* = 0 we get the classical rule, while for *s* = 1 we get the symmetric rule. We found that generalization performance increased with the parameter *s*, with the highest generalization at *s* = 1, again confirming that the symmetric rule might afford a representation that generalizes best.

To further assess the robustness of our results, we then investigated whether dropping the assumption of globally fixed values *α, β* would qualitatively alter the results. Again, this is motivated by the assumption that in biological agents, learning rules will not be perfectly static but might vary.

To capture this in the RL setting, first,we introduced noise to the parameters at every time step. That is, our update rule for the successor representation would become

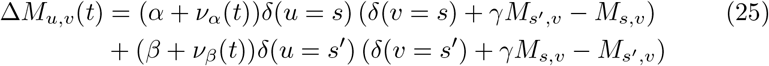

where *ν*_*α*_, *ν*_*β*_ is independent noise that is added at each timestep to the parameters. Secondly, we studied a condition where we introduced a state-dependent heterogeneity in the parameters. That is, for each possible transition *s, s*′ in the state space, there is a separate set of parameters *α*(*s, s*′), *β*(*s, s*′) which are randomly initialized at the start of learning and then fixed. The update rule thus is

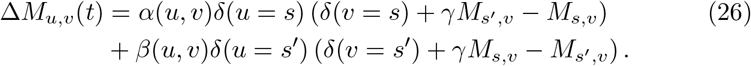

To determine the parameters, first some putative values 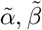 are drawn from Gaussian distributions with identical variance and different means 𝒩 (*µ*_*α*_, *σ*), 𝒩 (*µ*_*β*_, *σ*), and afterwards the absolute value of those is taken. This is to ensure that we don’t have negative parameters, which might result in qualitatively different and unstable learning rules. The means *µ*_*α*_, *µ*_*β*_ are then either set to (1, 0) to mimic the asymmetric with inhomogeneous parameters, or to 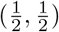 for the symmetric agent. In both variations we again find qualitatively similar results as before: even with noisy parameters, both agents are able to learn simple navigation tasks. The symmetric agent shows more variation in performance, but is in turn able to generalize better to new targets.

### 2.8 Maze tasks

The grid worlds we used to investigate the generalization capability so far were topologically very simple. In reality, biological agents will be exposed to more complicated navigation situations, with multiple paths and detours. We thus repeated our setup in a more contrived maze with multiple chambers and more convoluted paths. In particular, instead of keeping the environment static, when we tested the generalization abilities of the agents we introduced slight modifications, such that previously open paths were blocked. Again, in this setting the symmetric agent showed a better generalization, indicated by a higher probabilty of solving the new task with a number of steps close to the optimal one, as indicated by the distribution plots in Figure 9. However, in comparison to simpler grid worlds, both agents are worse at generalization in this setting, which might be caused by the additional complexity of the environment.

## 3 Discussion

Here, we have expanded the existing work on successor representation models of the hippocampus. We extended previous local learning rule models by including learning at two synapses, corresponding to CA3-CA3 recurrent synapses and CA3-CA1 feedforward synapses. In this model, we then studied the effect of making a local learning rule invariant to a reversal of time on the learned representations. We have found that the successor representations that are learned under such a learning rule correspond to encoding a transition structure which is also invariant to a time reversal - irrespective of the actual dynamics which are experienced. In particular, we could show that under such a symmetrized learning rule, place fields shift less when on a linear track, which is in line with empirical findings showing a distinction between place field shifting in CA3/CA1. Although we show that our model is able to replicate the observed shifting effects from Dong et al. (2021), further experimental data would be needed to corroborate these findings. Indeed, albeit CA3 place fields are generally reported as being more stable (Mizuseki et al., 2012), there are additional experimental and behavioural covariates to consider: For example, although Roth et al. (2012) find that the centre of mass of place fields in CA1 shift while those in CA3, are relatively stable, this holds only for familiar environments. In novel environments, both exhibit shift, although more pronounced in CA1. This speaks to the fact that a static, symmetric learning mechanism alone might be insufficient to account for all data, and plasticity might generally vary - both in strength as well as in symmetry. Additional experiments would be needed to disentangle such influences.

The local learning rule we have modified from Fang et al. (2023) is not the only one which could be used to obtain successor representations. Indeed, also George et al. (2022) and Bono et al. (2023) use local learning rules to learn these representations. It seems plausible that to obtain the successor representations we studied here, the exact shape of the learning rule is not important, as long as one can symmetrize it in an appropriate way. This might not even necessarily mean symmetrizing the plasticity kernel: For example, George et al. (2022) use STDP to learn feedforward connections between CA3 and CA1 neurons. Specifically, phase precession was used to provide a location code in the timescale of STDP, and importantly, in the absence of phase precession, a symmetric SR is learned in a biased random walk. Thus, it is plausible that precession in their model breaks the temporal invariance.

To understand the functional significance of such learning rules, we then went on to reinforcement learning experiments, where we trained an agent in navigation tasks. Here, we could show that a symmetrized learning rule affords a better generalization performance when the agent should navigate to a new target. This is interesting, because successor features have been explicitly employed in RL to obtain better generalization to new tasks (Dayan, 1993; Barreto et al., 2017, 2018). In particular, also when the SR was introduced as a model for hippocampus in the neuroscience literature, the generalization capability of such representations was measured (Stachenfeld et al., 2017; Gershman, 2018). In fact, it was argued in (Stachenfeld et al., 2017) that especially a successor representation that corresponds to a uniform policy should be beneficial for generalization. This was later also identified as a flaw of classical successor representation theory when linear reinforcement learning was suggested as a model for the hippocampal formation instead (Piray and Daw, 2021). In the latter, instead of storing the successor representation under the current policy, the representation under a default policy is stored, and the current policy can be efficiently represented by only encoding the deviations from default. This default policy in navigation is of course intuitively the uniform policy, which could be learned by an agent as soon as it encounters a new environment, in an explorative manner. Our symmetrized learning rule provides a middle ground between these two perspectives: the successor representation that one learns with this learning rule is depending on the current policy, but one can show that the reversible version is always closer to the SR of uniform policy than the SR under the original policy (see appendix). Thus, an agent does not necessarily have to start with exploration to construct a map of its environment, but can do so while performing a particular task. This is of course only possible due to the inductive bias that is inherent in the learning rule, which assumes that the state space is symmetric, since we observed that when this assumption is not true, the SR affords worse generalization.

Adapting learning mechanisms to symmetries of the data is a topic under ongoing investigation in biological and machine learning communities (Higgins et al., 2022; Mercatali et al., 2022). In reinforcement learning, symmetries are frequently considered under the framework of MDP homomorphisms (Van der Pol et al., 2020a; Mavor-Parker et al., 2022; Van der Pol et al., 2020b). This line of research for example aims at learning efficient abstractions of large state spaces into more amenable ones, or exploits known symmetries in the task structure to learn more efficiently. The symmetric state spaces we consider here are simple cases of an equivalence of states induced by the optimal value function: in a navigational problem in a metric space, any two states are equivalent with respect to the optimal value function if they have the same distance to the target (Givan et al., 2003). It would be interesting to investigate whether modifications of TD learning are also beneficial for the more general case of symmetries that are considered in this literature. Interestingly, it has been shown before that TD learning can be considered as performing a form of gradient descent if and only if the dynamics under the policy are reversible (Ollivier, 2018). This is intriguing, since TD-learning is known to be unstable in the continuous setting (Sutton and Barto, 2018) - if our symmetrized learning rule extends to this setting, then it might be possible that it could be useful for a more stable learning in symmetric settings.

Using a fixed learning rule for all synapses of a region might be a simplistic assumption, and in real networks, possibly a variety of learning rules are at play (Debanne et al., 1999). Our model of course does not fully capture this diversity, although we have investigated the stability of our RL model under noisy variations of the parameters. However, it would be easy to adapt the learning rules of individual neurons in such a sense that they have varying temporal profile. This could then possibly lead to a whole spectrum of successor features, each with its own sensitivity to future and past. In fact, it has been proposed before that the hippocampus encodes representation of both predecessors and successors (Namboodiri and Stuber, 2021), where representing preceding states has furthermore been suggested as useful for exploration in unsupervised reinforcement learning (Yu et al., 2023). These purely predictive respectively postdictive representations would be the two ends of a continuum of representations, with the symmetric learning rule in the middle. Thus, a model representing such a continuum could provide a more nuanced understanding of neural encoding and learning processes

Furthermore, it should be noted that there is a picture emerging where synaptic learning mechanisms are not but static but rather are controlled by neuromodulators, which could determine whether a certain firing pattern in pre- and postsynaptic neurons will result in potentiation or depression (Brzosko et al., 2019; Seol et al., 2007; Pawlak et al., 2010). Indeed, prominently (but not exclusively) dopamine and acetylcholine will modulate learning rules - both of which are important in reward-based learning and navigation (Brzosko et al., 2015; Sugisaki et al., 2011). In particular, it has been shown in a computational model that switching between plasticity regimes in a transmitter-dependent manner may result in successful spatial learning, in navigation tasks similar to the ones we considered here (Zannone et al., 2018). It may thus be an interesting avenue for future research to analyze such dynamically switching regimes in terms of successor representations to understand which predictive operation they compute. This might for example be achieved in the model presented here by modulating the parameters *α, β* depending on some external variable like reward.

Additionally, it is noteworthy that replay phenomena, where sequences of cells corresponding to recently visited states are reactivated in an orderly, time-compressed fashion, are taking place in hippocampus both in a forward as well as a reverse direction (Diba and Buzsáki, 2007; Foster and Wilson, 2006). From the perspective of reinforcement learning, for offline planning agents might indeed want to be able to simulate replay in the backward as well as the forward direction (Penny et al., 2013; Yu et al., 2021). These replay processes would provide yet another mechanism by which a symmetrized successor representation could be learned: Indeed, instead of modifying the TD learning rule during online learning, one could instead apply the classical learning rule during offline learning on replayed trajectories. This should again lead to a symmetrized representation, if forward and backward trajectories are replayed with equal probabilities. On the other hand, a symmetric learning rule in CA3 has been identified as a key mechanism to generate replay in the reverse direction (Haga and Fukai, 2018).

Our model naturally is a broad oversimplification of matters and lacks biological realism. This is not a problem per se, because we aimed here at analytical amenability and to expand on the theory of successor representations, which operates on the computational level. Still, our focus was obtaining an explicit relation to successor representations by considering CA3/CA1 in isolation, with external input synapses not subject themselves to learning. In reality, it is now accepted that there is not a simple forward pass through the hippocampus, but rather there are projections from the deep layers of entorhinal cortex back into the superficial layers (which provide input to hippocampus), essentially creating a loop (Kumaran and McClelland, 2012; Canto et al., 2008). With plasticity also happening at these synapses, one would then obtain a model that is not as simple anymore as the one we presented here. Studying the representations in such an extended model, which would include learning for example in synapses from HC to EC, and whether these can still be framed in Successor representation theory, would be an interesting future research direction and could possibly build a bridge to computational models which include hippocampal-entorhinal interactions (George et al., 2023a; Whittington et al., 2020). Speculatively, when including spatially selective cell types like grid cells from the entorhinal cortex in the loop, the general result that HC represents successor features should not change. Indeed, taking the input from multiple grid cell modules together, these cells provide a unique encoding of spatial position (Fiete et al., 2008). This is thus an injective function of the current spatial state, just as we used as input for our system. These coordinates, as they encode space, naturally benefit from a symmetrized representation in CA3. One step further one could then include additional, non-spatial input externally to CA1. This would then result in a successor representation of the non-spatial features, conditioned on the spatial features. That is, a long-term prediction of expected stimuli, given current position in space. Connections back from hippocampus to EC might in turn be included to plasticity rules of a similar spirit as the one we used here. Spatial cell types like grid and head direction cells are widely believed to follow specific connectivity implementing a continuous attractor (Zhang, 1996; Samsonovich and McNaughton, 1997; Khona and Fiete, 2022). Indeed, typical continuous attractor models are characterized by their fixed connectivity patterns, establishing an attracting manifold as well as a velocity modulated update mechanism thereon. Projections from CA1 to EC would then putatively implement a predictive mechanism that would be focussed on predicting the outcome of this rigid update mechanism using the predictive representation of the conjunctive spatial and non spatial features obtained from the hippocampal loop. However, as this is a slightly different kind of model, there might not be a straightforward result linking the weights to a successor representation type quantity, but this would be an interesting avenue for further computational work.

Finally, we want to mention that the hippocampal formation is also of high interest in studying generalization in human cognitive neuroscience (Theves et al., 2021; Garvert et al., 2023). Since evidence is growing that the systems which are partaking in spatial representations are also recruited to encode more abstract variables, possibly forming ‘conceptual spaces’ (Nitsch et al., 2023; Bottini and Doeller, 2020; Constantinescu et al., 2016), it might be interesting to understand whether an inherent bias for symmetry also shapes these representations. In particular, one might speculate for which domains of cognition such a bias for symmetry or a metric space structure might be adequate, and when it would not be. This might for example be tested by constructing explicit tasks where dynamics are reversible and compare subjects accuracy in predictions to such tasks where the dynamics are not.

In conclusion, our model contributes to the theoretical framework of hippocampal predictive representations, both on the mechanistic level through suggesting the use of symmetric local learning rules, as well as on the functional level, where such learning might be useful for generalization in spatial learning.

## 4 Methods and Materials

## 5 Methods

### 5.1 Neural Network Model

We consider two populations of rate-based neurons *p*^1^ ∈ ℝ ^*m*^, *p*^2^ ∈ ℝ ^*n*^ respectively. The population *p*^1^ is recurrently connected via a matrix of synaptic weights *W*^*r*^ ∈ ℝ ^*m*×*m*^, and feeds its activity forward to population *p*^2^ via the matrix of synaptic weights *W*^*f*^ ∈ ℝ ^*n*×*m*^. Both regions receive additional external inputs, *ϕ*^1^, *ϕ*^2^, and decay to equilibrium in the absence input. The temporal evolution of the populations is then modelled by the following differential equations:

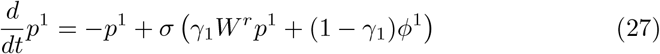

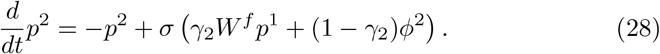

Here, *σ* is an activation function, and the *γ* parameters control the relative weight of the different types of inputs. On the computational level, these parameters will correspond to the discounting factor of the successor representation, effectively scaling how far in the future predictions are made.

On the level of biological implementation, there are two speculative interpretations one may have for these parameters. First, the *γ*_*i*_ might be associated with acetylcholine. Indeed, this transmitter is believed to control the relative strength of external and internal input in hippocampus (Hasselmo, 2006). In particular, a low value of *γ* would correspond to an encoding mode, where the network is relatively stronger driven by external input and thus susceptible to encoding associations or transition structures between these inputs. For high *γ*, the network is in retrieval mode and its outputs will depend more on the recurrent weights and therefore previously learned transition structure. Indeed, this interpretation has been pursued in (Fang et al., 2023), who showed how a recurrent net could learn weights at low *γ* and then use those learned weights to retrieve successor representations at arbitrary high *γ*. Another interpretation of *γ* would be that it represents the respective proportions of inputs of different kinds a certain subpopulation receives due to anatomic difference. That is, it could for example encode the number of synaptic contacts from other CA3 axons for a certain subpopulation - then understood not in absolute terms, but compared to other cells of the same type. Indeed, there is evidence that both connections in as well as connections to hippocampus are not homogeneous but vary along the septotemporal and proximodistal axis (Ishizuka et al., 1990; Witter, 2007). Thus, it might be plausible to assume a varying degree of strength of the different input types. In this work, however, we do not explicitly model these biological details, and thus the precise role of *γ* remains undetermined.

Assuming the weights (and the external inputs) change on a slower timescale than the population dynamics, we can let the above differential equations go to equilibrium for analysis - in practice, we can also integrate the ODE above with a timescale *τ* orders of magnitude smaller than the timescale of learning. If *σ* is approximately linear, this results in

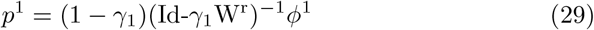

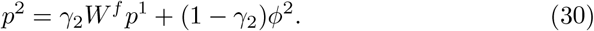

We assume that the animal receives inputs depending on the current state of the environment, which we will denote by the process *S*_*t*_. In the reinforcement learning setting, the agent influences the way in which the state-process is sampled by selecting actions *A*_*t*_, but at this point this is not relevant since we are only building up a predictive representation of states, not actions. The inputs to the two neural populations thus take the form

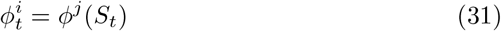

where *ϕ*^*i*^ is a function mapping the state space to an activation pattern in neural space. If the state space is discrete, we can also write this as

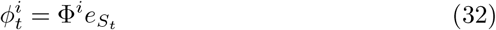

with Φ^*i*^ now being a matrix of appropriate dimension, and 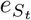 is the unit-vector corresponding to state *S*_*t*_, which hence selects the corresponding activation pattern from Φ^*i*^s columns. In the special case Φ^*i*^ = Id, we have a so called one-hot encoding, where hence every cell corresponds to a distinct state and fires if and only if the agent is in that state.

#### 5.1.1 Learning Rule

The synaptic weights - both feedforward and recurrent - in our model are subject to a learning rule which is controlled by two parameters *α, β* ∈ ℝ:

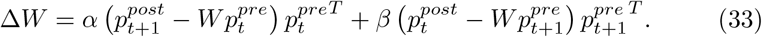

Here, *p*^*pre*^, *p*^*post*^ denote the vectors of pre- and postsynaptic activities. The parameters *α, β* do not model a biological substrate, they control the qualitative behaviour of the learning rule: they serve as a (crude) discrete approximation for a continuous plasticity kernel as is typically observed in spike-time dependent plasticity protocols. Indeed, for *α* = 1 the learning rule strengthens connections where post-synaptic activity at the later timestep was high (measured relative to overall input), and presynaptic activity at the earlier timestep was high as well. In contrast, for *β* = 1, the learning rule strengthens connections where postsynaptic activity at an earlier timestep was high, and so was presynaptic activity at the later timestep.

Note that a synaptic update of the form

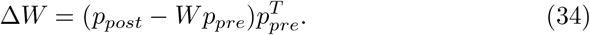

may be seen as performing a sort of conditional expectation or regression objective: At equilibrium, the expected update in synaptic weights, given the current activity, should be zero

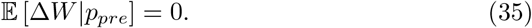

We have that this holds in general only if

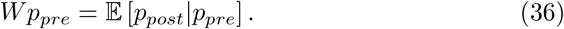

That is, if *W* encodes an optimal prediction of the post-synaptic activity given the presynaptic activity. Hence, heuristically, *α* emphasizes to learn weights that are optimally tuned to predict the next state of their output. In our setting, the cells are driven by external input identifying the state *S*_*t*_ of the world, so this amounts to optimally predicting the next state *S*_*t*+1_. In turn, *β* emphasizes a backward prediction, where *S*_*t*_ should be predicted by *S*_*t*+1_.

### 5.2 Forward and Backward Process, Reversibility

Recall that a time-homogeneous Markov process on a finite state space 𝒮is determined by its one-step transition probabilities

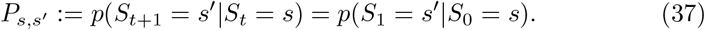

Under standard assumptions, the process has a unique stationary distribution (Norris, 1998; Seabrook and Wiskott, 2023), that is, a probability distribution *π* over states such that

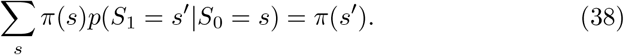

That is, if we are in stationary distribution, the probability mass is left invariant when transitioning according to our process. In particular, if we assume we start in stationary distribution, then *p*(*S*_*t*_ = *s*) = *π*(*s*) ∀ *t*. We may now define the reverse process of our original process, by simply reversing the order of time. That is, define a process *R*_*t*_ with transition probabilities

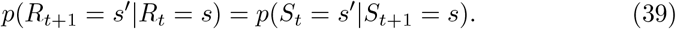

These probabilities will generally depend on the timestep *t*, but they will not if our process *S*_*t*_ is in stationary distribution. In that case, we again have a single matrix that encodes all transition probabilities:

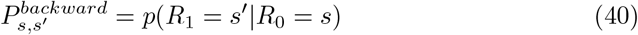

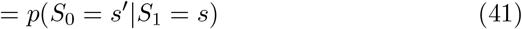

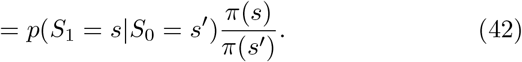

In matrix notation, we have

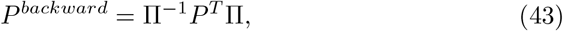

where Π is a diagonal matrix with the values of *π* on the diagonal. It is easy to see from this definition that *P*^*backward*^ is again a valid transition matrix, and that it has the same stationary distribution as *P* = *P*^*forward*^.

Forward and backward transition probabilities are equal if and only if the process fullfills the *detailed balance* condition

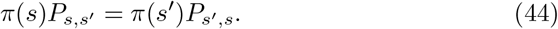

In that case, the process is called reversible. In Equation 19 we have used a weighted sum of transition probabilities of the forward and backward process Note that the symmetrized process with transition probabilities

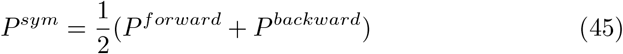

will always be reversible, as can either be seen by a simple algebraic calculation, or by the following argument: Let *P* ^∗^ be the *adjoint* of *P* with respect to the inner product induced by *π*. This means that for any vectors *x, y* we have

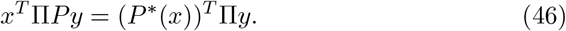

We then have that

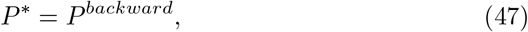

as may be seen by plugging in the definition of *P*^*backward*^ into Equation 46. For the adjoint, we have that (*P* ^∗^)^∗^ = *P*. Thus, in particular,

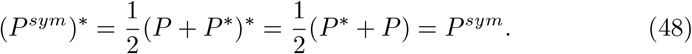

This also explains that it is warranted to talk about a ‘symmetrization’ here: indeed, the adjoint with respect to the standard inner product is the transpose of a matrix *A*, in which case one would obtain the standard symmetric part of a matrix, that is, 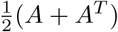.

### 5.3 Reinforcement Learning and Successor Features

In Reinforcement learning, one generally considers a Markov decision process (MDP), that is a tuple (𝒮, 𝒜, *T, R, γ*), where 𝒮, 𝒜 are the sets of possible states and actions, *T* (*s*′ | *s, a*) gives the transition probability from state *s* to state *s*′ when choosing action *a, R*(*s, a*) is obtained reward when chosing action *a* in state *s*, and *γ* is a discount factor. The goal in RL usually is to find an optimal policy, that is a probability distribution *π*(*a*| *s*) over actions, given states, which maximizes the expected discounted cumulative reward

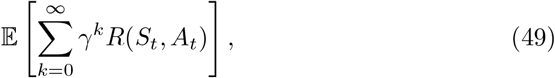

where the states and actions *S*_*t*_, *A*_*t*_ are sampled according to the policy *π* and the transition probabilities *T*. In our experiments, we are only interested in the particularly simple case where the transitions are deterministic - that is, taking an action *a* in state *s* surely leads to a specified state *s*′(*a*). Furthermore, we only consider the situation where the reward is a function of the state only. However, all definitions in the paragraph below readily generalize to functions of states and actions. Assume a fixed policy *π* and a process *S*_*t*_ following said policy. Now consider any mapping from the state space

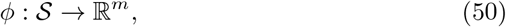

which we can interpret as an observable generated by the state space. Then define a function on the state space

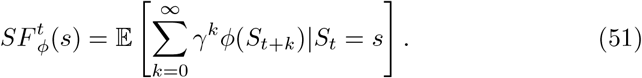

*SF*_*ϕ*_ thus gives the expected (exponentially weighted) cumulative future sum of the observation or feature *ϕ*, given the current state *s*. This is hence called a ‘successor feature’ in the literature. Define 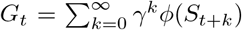. We then have

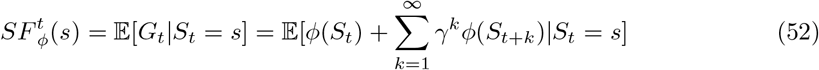

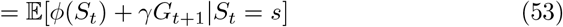

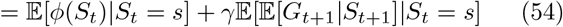

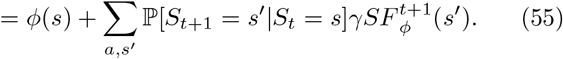

In particular, if the transition probabilities of *S*_*t*_ are time-homogeneous, we see that *SF* itself does not depend on *t*, and we can write

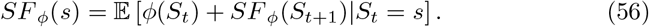

Temporal Difference learning (TD-learning) uses this relationship to construct a target to update an estimate of *SF*_*ϕ*_ online. Indeed, if 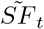 is the current estimate the agent has for the successor features, then consider update rules

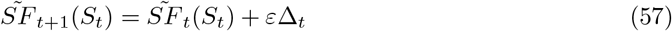

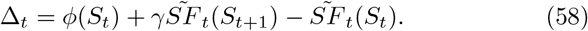

Then we see that if 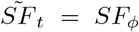, we get 𝔼 [Δ_*t*_|*S*_*t*_ = *s*] = 0. Thus, here 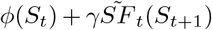 is used as a bootstrapping estimate of the target 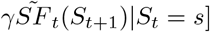, and 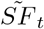 is updated according to the error to that target - in total, Δ_*t*_ is thus also called TD-error.

Successor features encompass important special case examples: For the choice of *ϕ* = *R* the reward function, *SF*_*R*_ becomes the value function: The value function under a policy *π*, which is typically denoted as *V* ^*π*^, hence encodes the weighted cumulative sum of expected rewards, given the current state. In particular, the current estimate of the reward function may be used to define a new policy as

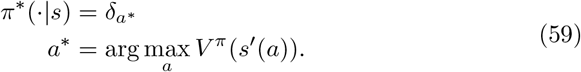

That is, the policy deterministically selects the action *a*^∗^ which leads to the next state with the highest value according to the current value function estimate. Iterating this process then successively learns a better value function - this process of updating an estimate of the value function and then choosing an optimal policy with respect to it is the basis of the classical TD-learning algorithm (Sutton and Barto, 2018). Indeed, for our navigation experiments we use this approach, only replacing the maximum in Equation 59 by a softmax which smoothens the transition probabilities. Note that the above assumes a model of which actions lead to which next states - which posing as given should be a sensible assumption in navigation problems - but a completely model-free approach simply computes the value of a state-action pair instead.

#### Successor Representations

In the case that we have that *ϕ* is in fact an injective mapping, we can define a modified version of successor features as

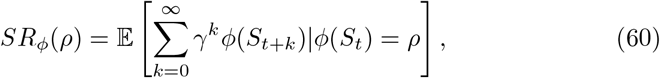

where *ρ* ∈ *ϕ* (𝒮). Here, the injectivity of the mapping is necessary to ensure *SR* defined as above still enjoys desirable properties like homogenity in time and the Bellman equation, but besides of that one could also define a similar quantity without using injectivity. Now, in the injective situation, we would like to call the above ‘successor representation’. This is because when we take mapping *ϕ*(*s*) = *e*_*s*_, where *e*_*S*_ is a unit vector in ℝ^|*S*|^, then we obtain

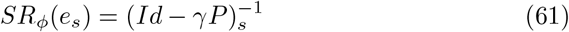

which corresponds to the classical definition of successor representation. In general, if we assume Φ ∈ ℝ^*m*×|*S*|^ is the matrix of features (could also be an operator if we allow for continuous state space), then we have that

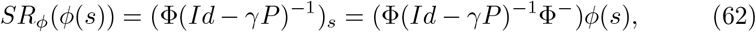

where Φ^−^ is such that Φ^−^Φ = Id_|*S*|_.

#### Generalization and Successor Features

The idea behind using successor features for some set of function *ϕ*^*i*^, instead of simply directly computing the the value function is that of generalization/transfer: Assume the reward function *R* can be written as a linear combination of the features *ϕ*, that is

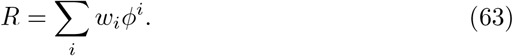

Then also for the respective predictive representations one has

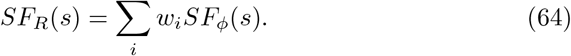

Now assuming that the reward changes to a new function 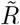, which also can be expressed with the features, then the only thing that has to be relearned is the weights *w*_*i*_, while the successor features *SF*_*ϕ*_ can be reused. Thus, one may then generalize more easily to a new task, since the transition structure under a policy is essentially already learnt. In particular, in a discrete state space, a fixed reward function can *R* can of course be encoded by a reward vector **R**, that is

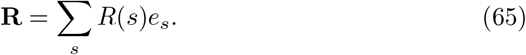

The value function is then simply given through the classical SR as

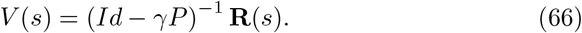

Thus, for a fixed policy, this separates the computation of value into learning a successor representation and learning a reward vector. Our navigation experiments probe the generalization capability of this approach, by introducing a new reward vector 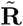, after an optimal policy and a SR for a reward vector **R** have been learned. Importantly, only the reward vector is allowed to be relearned, while the SR has to be reused from the previous task.

#### Symmetric TD-learning

In our reinforcement learning experiments, we use a modified version of TD-learning to mimic the behaviour of the local learning rule in the SR-network. In practice, Successor features are typically parametrized by some parameter *θ* (e.g., the weights of a neural network), which is then updated to reduce the TD-error. That is, for each value of *θ* we obtain a map 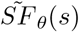 of states. We can then update the parameter *θ* via

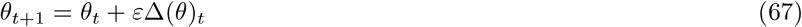

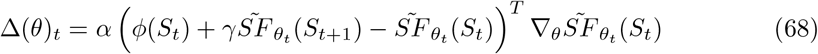

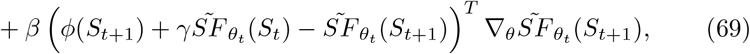

where *α, β* are parameters corresponding to the ones controlling the local learning rule. In particular, for *α* = 1, *β* = 0 one obtains the classical TD-learning rule for function approximation (Sutton and Barto, 2018) and for *α* = 0, *β* = 1 one obtains the ‘predecessor representation’ (Yu et al., 2023). The case *α* = *β* yields a symmetrized version of TD-learning. In our experiments, we use a particularly simple version of the above: for a discrete state space, one can simply parametrize 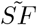 by means of a matrix *M* ∈ℝ^*d*×*k*^, where *d* is the number of features and *k* is the number of states. Then one has 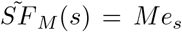, and hence the update rule

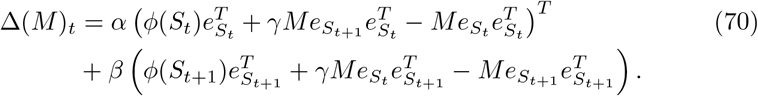

### 5.4 Successor Representations and Hitting Times

In the theory of Markov chains, it is a common exercise to study the fundamental matrix of the Markov process, which encodes properties about the process via the first hitting times of states (Levin and Peres, 2017). The first hitting time of a state, 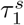, is defined as the first time a process hits the state *s*. The successor representation may similarly be interpreted as an operator that encodes certain expectations related to the first hitting times. Indeed, in subsection G.1 we prove the following formula:

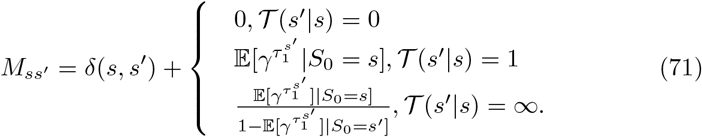

Here, 𝒯 (*s*′ | *s*) denotes the largest number of times which state *s*′ may be hit with nonzero probability when starting from state *s*. It may only take values 0, if *s*′ may not be hit from *s* at all, 1, if *s*′ is a transient state, or ∞, when the process may arbitrarily often return to a state. Equation 71 provides a convenient reformulation of the value function. In particular, when we are in the setting of a navigation task, where the goal is navigating to target *s*^∗^, the reward is a unit reward at *s*^∗^. Hence, for an optimal policy *π*^∗^ which only chooses among shortest paths to the target, the successor representation becomes

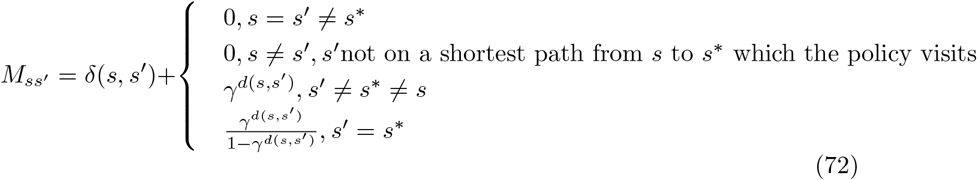

where *d*(*s, s*′) is the distance on the state space. The value function may then be read of as the column corresponding to *s*^∗^. We use this formula in subsection G.1 to prove a stability result of optimal policies in the navigational setting under symmetrization: If we are in the deterministic navigation setting, and our policy *π* is optimal, then after taking the symmetrization of the transition probabilities and computing the value function with these probabilities, the policy *π* is still optimal.

### 5.5 A predictive representation leads to backward shift only in the asymmetric case

Here we want to give an analytic explanation for the stronger backward shifts of the centre of mass observed in Figure 4 when using an asymmetric learning rule, while a symmetric learning rule does not result in a shift. Recall that we simulated an agent repeatedly running on a linear track, always in the same direction, which was then reset to the starting position. The effect is most easily analyzed if we ignore boundaries and go to a continuous situation. Let’s thus assume an agent is repeatedly running on the real line R. In the beginning, before learning, the cells in our model are just driven by the external input, and thus can be taken equal to the features. That is, cell *j* will take the value *ϕ*^*j*^(*x*) when at position *x*. After learning, it will instead encode a successor feature. With continuous space it is also more convenient to assume continuous time.

**Figure 4:**
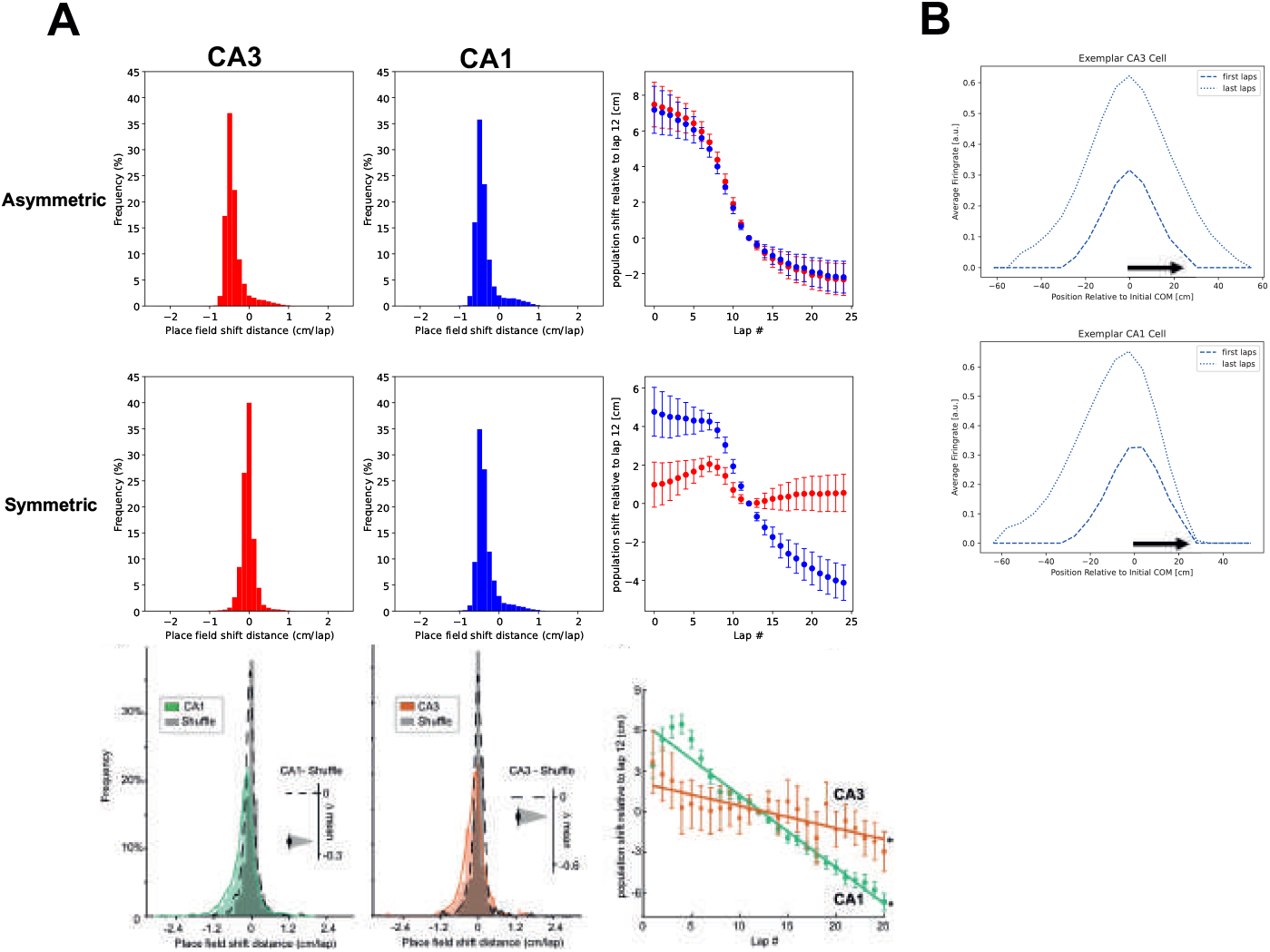
Symmetric learning rule leads to more stable place fields in linear track. We simulated an experiment with a rat repeatedly running on a linear track, similar to Dong et al. (2021). A two-layer SR network was used where the recurrent weights had a symmetric (**A, middle row**) or asymmetric (**A, top row**) learning rule. In the the symmetric case, there is less shift of the centre of mass of place fields in the modelled CA3 population (*red*) than in the CA1 population (*blue*), which is not the case in the asymmetric version. Histograms show distribution of shifts comparing last five laps versus first five laps, while the rightmost plot shows shift relative to the 12-th lap. The results in the symmetric case are qualitatively similar to data (**A, bottom row**) from *Ca*^2+^ recordings of hippocampal neurons in a similar experiment - figure adapted from Dong, C., Madar, A. D., & Sheffield, M. E. (2021). In **B**, we show firing rates of an exemplar cell from CA3 and CA1 respectively, where the symmetric learning rule is used for CA3. The firing rates in each position are averaged over the first and last five laps, and plotted relative to the centre of mass in the first laps. With experience, only the place field in CA1, not the place field in CA3 shifts backwards (arrow indicates direction of travel).

**Figure 5:**
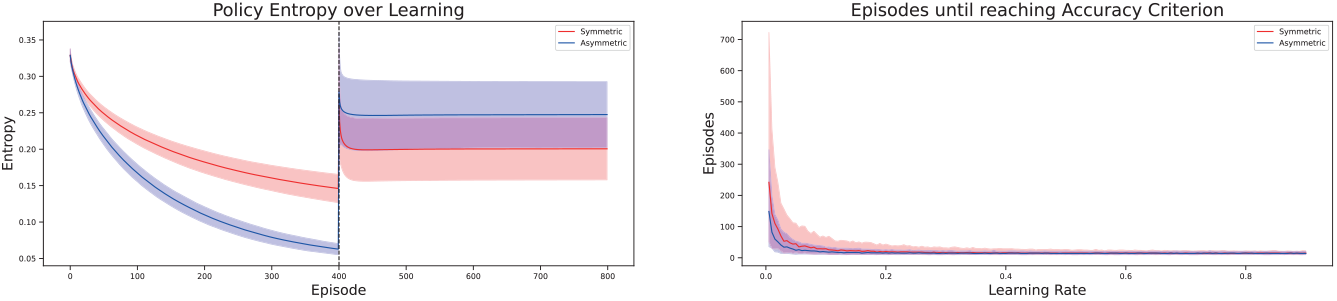
Comparison of policy entropy and sensitivity to learning rate among agents with symmetric and asymmetric learning rule. **Top:** Symmetric agent shows more sensitivity to learning rate parameter for lower learning rates. We trained the agents repeatedly until a fixed accuracy in navigation to the target was met. We then recorded the number of episodes it took until that criterion was reached. Curves show median and interquartile range of this number for the two agents. **Bottom:** Policy entropy of the two agents show different trajectories in the generalization experiment. We calculated the entropy of the agent’s policy, averaged over all states, at the end of each episode. This reveals that during learning to navigate to the first target, the symmetric agent has more entropy, which is then reversed when the new target has to be reached.

The successor representation may be easily transferred to this setting (George et al., 2023b), where the definition then is

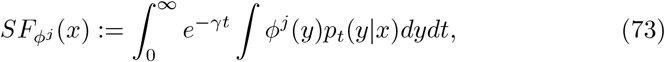

where *p*_*t*_(*y*| *x*) is the probability of transitioning from state *x* to state *y* in time *t*. The centre of mass (COM) of a cells’ firing field is just the spatial mean of the cells’ activity when the latter is normalized to be a probability distribution. That is, for example before learning, when the cell *j* fires at position *x* according to the feature *ϕ*^*j*^, the *j*-th centre of mass is

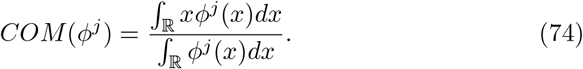

To understand the observed shifting effect in *COM*, it is now instructive to study the successor feature with deterministic dynamics: Assume we are moving with constant velocity *v*, that is,

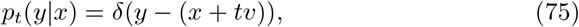

then the successor feature takes the form

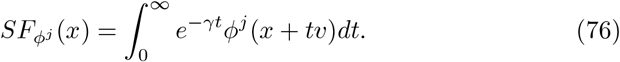

Now, taking the mean over all positions yields:

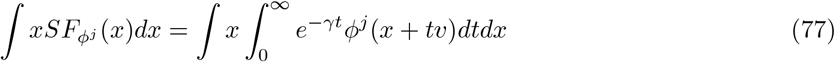

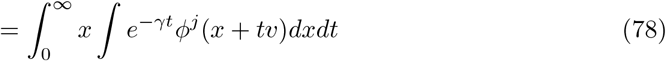

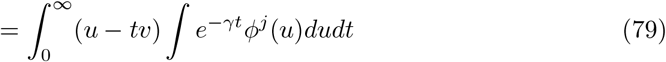

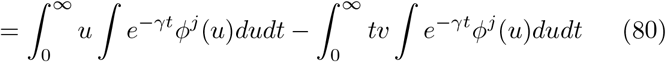

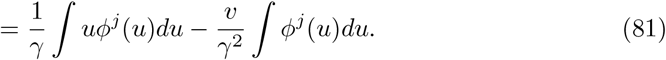

Similarly we have

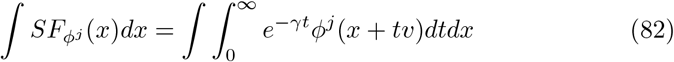

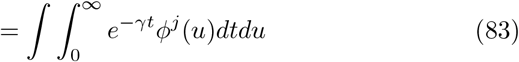

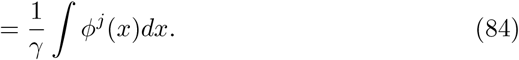

Thus, in total we obtain

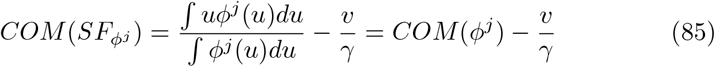

That is, the center of mass is shifted backward by a factor which is controlled by the prediction timescale *γ* and the velocity of the movement. We thereby also obtain the seemingly new prediction that faster running speed and smaller terminal place field size (as a proxy for *γ*) should result in bigger shifts. Now if our forward process is a deterministic process with constant velocity −*v*, then conceptually, our backward process is also deterministic with velocity *v* (although stricly speaking, there is no stationary distribution). In particular, the symmetrized process is a process which in an infinitesimal amount of time travels either forward with velocity *v* or backward with velocity *v* with equal probability. The shifts thus cancel and the centre of mass stays the same as for the initial feature. The same relation of shift and velocity/timescale holds also when the dynamics come from a Brownian motion with constant drift, that is 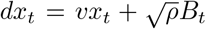, as we show in appendix B. Now when we symmetrize the Brownian motion with drift, the drift term will disappear, that is, we have a standard Brownian motion. It is thus clear that under this motion, there should again be no shift of the centre of mass.

### 5.6 Experiments

### 5.7 Convergence Experiments

For our initial convergence experiments (Figure 3, top row) we simulate our model in a circular random walk setting with 30 states and use *n* = 40 cells in each layer. The input at each state is drawn from an i.i.d. Gaussian distribution (*σ* = 0.1) (i.e., for each cell and each state we draw a value from a Gaussian distribution, this is then assumed to be the input for that cell whenever the specific state is visited). Throughout this and the following experiments, we use a moderately high value of *γ* = 0.7 - meaning a relatively large timescale/small discounting. This is an arbitrary choice, but relatively high values have been used in the past in SR-theories of hippocampus (Geerts et al., 2020).

We observe 100000 transitions, and repeat the experiment 30 times. We then repeat the same experiment (top row, right of Figure 3), also drawing the transition matrix *P* randomly. To track the convergences, we consider the loss terms

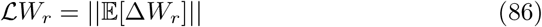

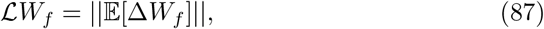

where the terms on the right hand vanish at convergence and are defined in Equation 139. The resulting convergence curves are shown in Figure 3. These experiments are conducted with a linear activation function, we also repeat this experiment with the activation functions **tanh, relu** as shown in Figure H.1

To check the effect of different values *α, β*, we conduct a parameter sweep in a circular random walk, and random Gaussian features as above. For each combination *α, β* we run 30 random initializations for 1000 iterations, and check this combination once for the recurrent layer and once for the feedforward layer. Here, we track convergence similar as above, but now we take the fraction

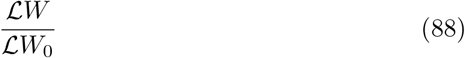

with *W*_0_ the initial weight, to judge if we are moving towards or away from equilibrium. This results in the matrices shown in *Figure* 3. To speed up the experiments, we use a smaller network for these experiments, with 20 neurons in each layer and 10 states. Again we use *dt* = 0.1.

### 5.8 Linear Track Experiments

To simulate the animal running on a linear track, we partitioned a track of length 300*cm* in 50 discrete states and let the agent perform a rightward biased walk that either took a step to the right (*p* = 0.9) or stayed in place (*p* = 0.1). We chose a time step of *dt* = 0.4, corresponding to a velocity of ≈ 15*cm/s*. The agent would run 25 laps on the linear track, with a short resting phase in between two laps. We set up a two-layer model, with *n* = 100 cells in each layer, and again *γ* = 0.7. For the feedforward weights, we always used the parameters *α* = 1, *β* = 0, while for the recurrent weights we tested both the symmetric case 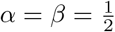 as well as the asymmetric case *α* = 1, *β* = 0. We then performed an analysis approach as in Dong et al. (2021). For each lap, we collected the mean activities of each layer per state and computed the center of mass (COM) by the formula

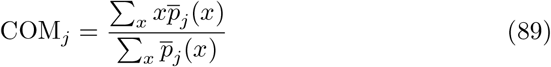

where *x* is position on the track and *p*_*j*_ (*x*) denotes the mean activity of cell *j* when at that location. To obtain a distribution of the overall shifts of the COMs over all laps, we calculated the average COMs for the first and the last five laps and subtracted them from each other, obtaining the histograms in Figure 4. To track the evolution of the shift, we subtracted the COMs from the 12-th lap, which yields a more gradual tracking of shifts. We plot the results of this approach in the scatterplots in Figure 4.

### 5.9 Reinforcement Learning

We investigated the generalization capabilities of a symmetric over an asymmetric learning rule in TD-learning in different scenarios.

#### 5.9.1 Navigation Experiments

Since the hippocampal formation is prominently involved in navigation tasks, we first focused on tasks of such nature. We thus studied different variations of the same general problem setup: Given a grid environment with a deterministic transition structure, the agent would receive a unit reward only when arriving at the designated target state *s*_*T*_, upon which the episode is terminated, and the next episode starts in an initial state *s*_0_, drawn uniformly at random. The agent is thus encouraged to navigate from all possible starting states to the target state to receive a reward. We used the Neuronav toolbox (Juliani et al., 2022) to implement the grid environment and implemented our modified SR-agent in the framework of that toolbox. For completeness, let us here again give a concise summary of how the agent learns. The agent possesses a world-model *P* (*s*′| *s, a*) and updates an internal estimate *M* of the successor representation as well as an estimate *w* of the reward vector. Having observed a transition from state *s* to state *s*′ via action *a* and a reward *r*, the updates are

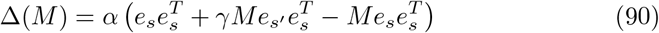

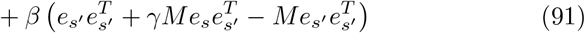

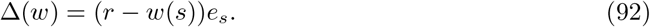

In summary, this is how the agent selects actions and updates its internal estimates:

- Given the current state *s* and current estimate *M* for the successor representation, compute *q*(*s, a*) = _*s*_′, _*s*_′′ *w*(*s*0′′)*M* (*s*′′ |*s*′)*P* (*s*′ |*s, a*)
- Select an action according to *π*(*a*|*s*) = softmax(*q*(*s, a*))
- Observe next state *s*′ and reward *r*
- Update reward estimate *w* and successor representation estimate *M*.

To check for generalization capability, we would train agents which utilize the respective learning rules (symmetric,asymmetric) to navigate to a fixed *s*_*T*_ first. We trained the agent either for a fixed number of episodes (we used 400 episodes with a maximum number of steps of 400), or until a fixed accuracy criterion was met (the mean deviation from optimal performance for the preceding 8 episodes was lesser than 2). Then, we would randomly select a new target state 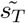 from among all possible states. Importantly, after the modification of the target states, the agents were only allowed to modify the reward-prediction vector *w*, but not the successor representations *M* themselves. That is, they could only learn the reward structure of the new task, but had to rely on the transition structure that was encoded during learning the previous task, which also depends on the policy. We repeated this experiment for 200 times, with randomly drawn combinations 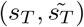 in each repetition. The results of these experiments are depicted in Figure 6. For the first experiments reported in the main text, we used a learning rate of 0.1 and set *γ* = 0.7.We chose a relatively high value of *γ*, as this is typically done when modelling hippocampal place cells in standard navigation tasks (Stachenfeld et al., 2017; De Cothi and Barry, 2020), but we repeated the second kind of analysis for different values of *γ* and the learning rate, as shown in Figure H.7.

**Figure 6:**
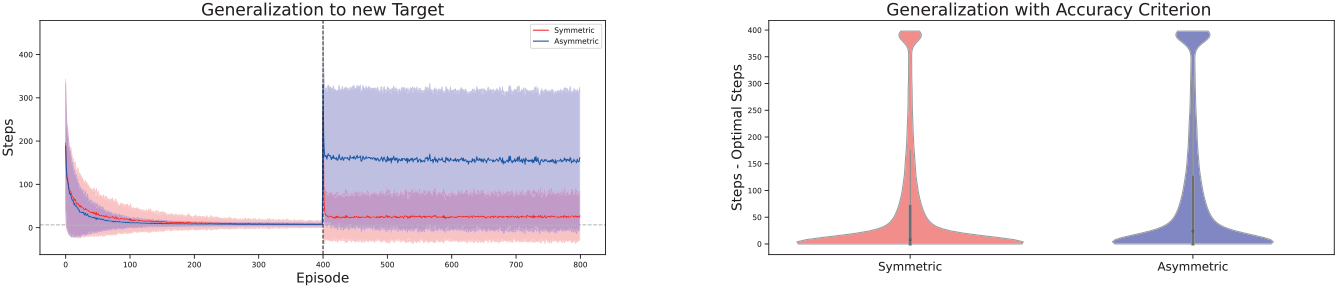
Symmetric successor representation agent affords better generalization in simple navigation tasks. **Top:** Agents started in random locations in the environments and had to learn to navigate to fixed targets. After 400 episodes, reward location was switched to a new random location, where agents could only relearn the reward prediction vector but not the SR. (Generalization) performance is visualized by total number of steps taken per episode, for an agent using the classical rule (blue) and an agent using the symmetric rule (red). Dashed line indicates change of target location. We show the average performance over different environments as performance is qualitatively similar, see Figure H.3 for plots in individual environments. **Bottom:** Similar to left plot, but instead of switching target after a fixed number of episodes, the target was switched when the previous target was found with a fixed accuracy. Violin plots show distribution of suboptimality (steps - optimal number of steps) over all environments, for individual environments see Figure H.4. For an outline of the environments see Figure H.2

**Figure 7:**
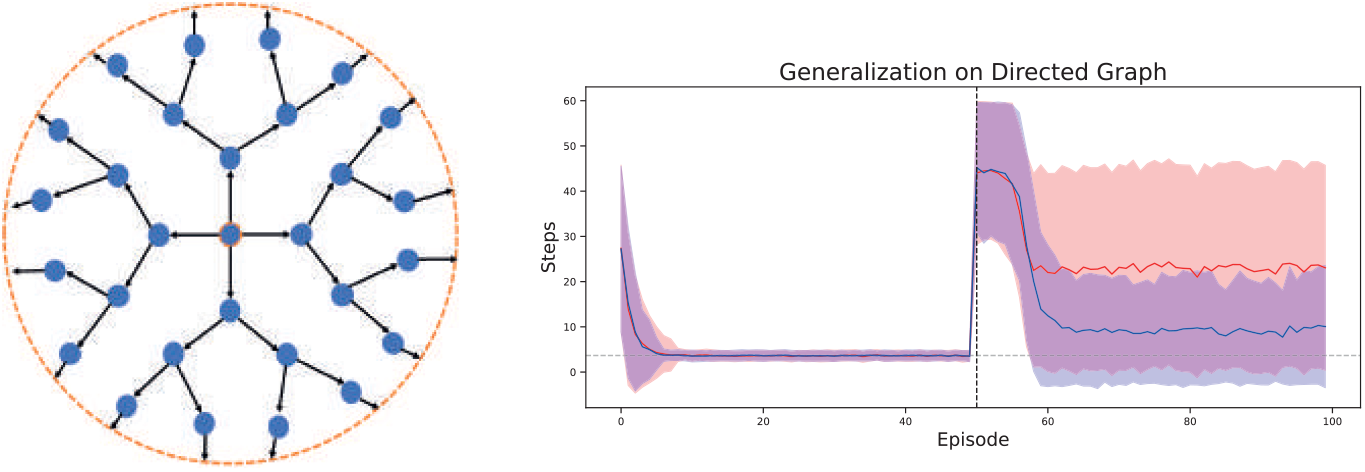
Symmetric learning rule provides no advantage in generalization experiment on a directed graph. We conducted the same kind of experiment as in Figure 6 on a directed graph. **Left**: The state space is tree-like, with the addition that from the leaf nodes at the last level one travels back to the central node (orange dashed line). **Right**: In this scenario an SR agent with the classical learning rule (blue) performs better in generalization than one with the symmetric learning rule (red).

**Figure 8:**
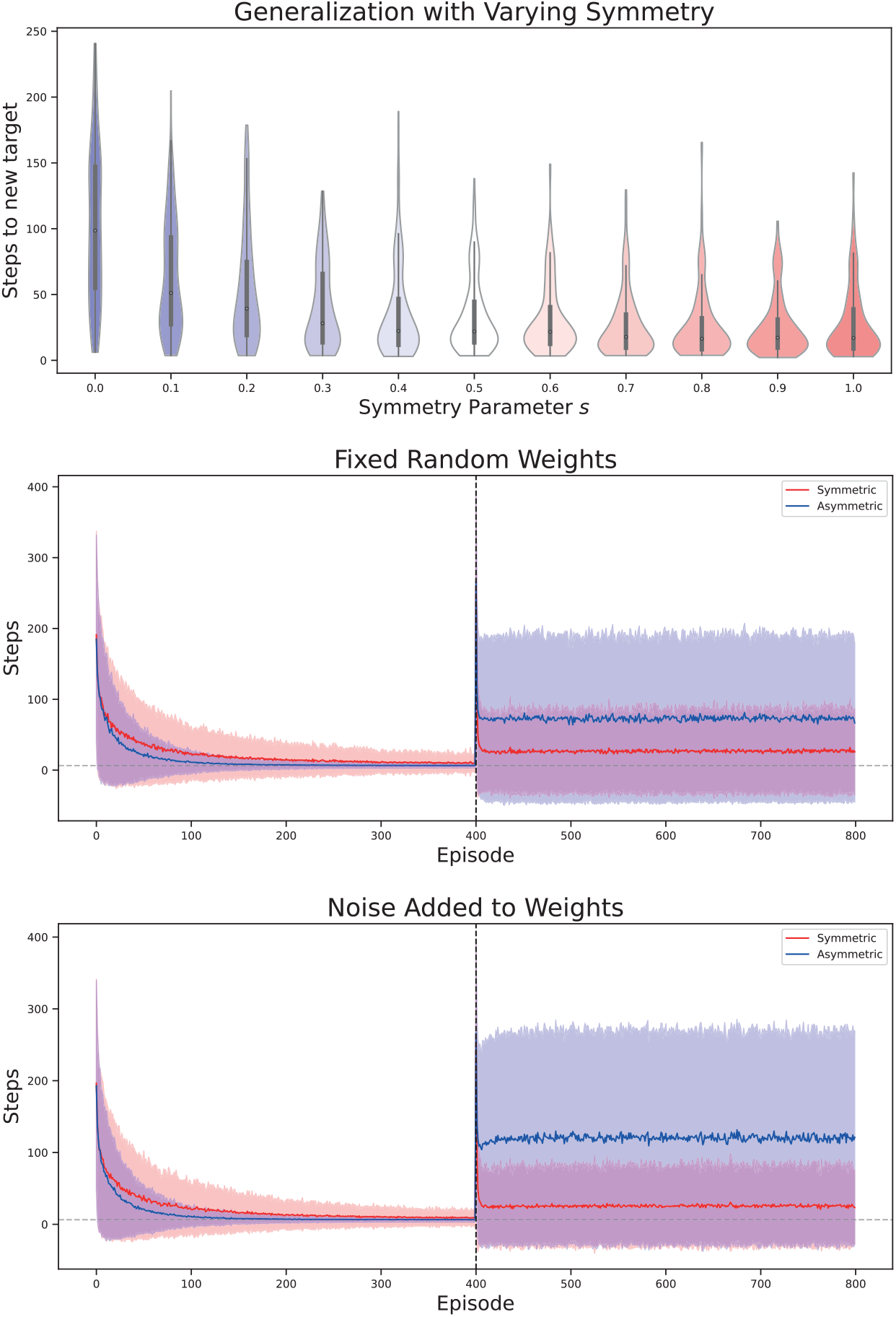
Variations of symmetry in the learning rule. Experiment for all plots is the same as in Figure 6. **Top:** Generalization for parameters 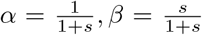. Violin plots show distribution of differences (steps-optimal number of steps) when evaluated on new targets. Distributions broaden towards the optimal value of 0 with increased symmetry. **Middle**: Generalization with noise added to *α, β* at each timestep. **Bottom:** Generalization with parameters *α, β* randomly initalized for each pair of states. Figure H.2

**Figure 9:**
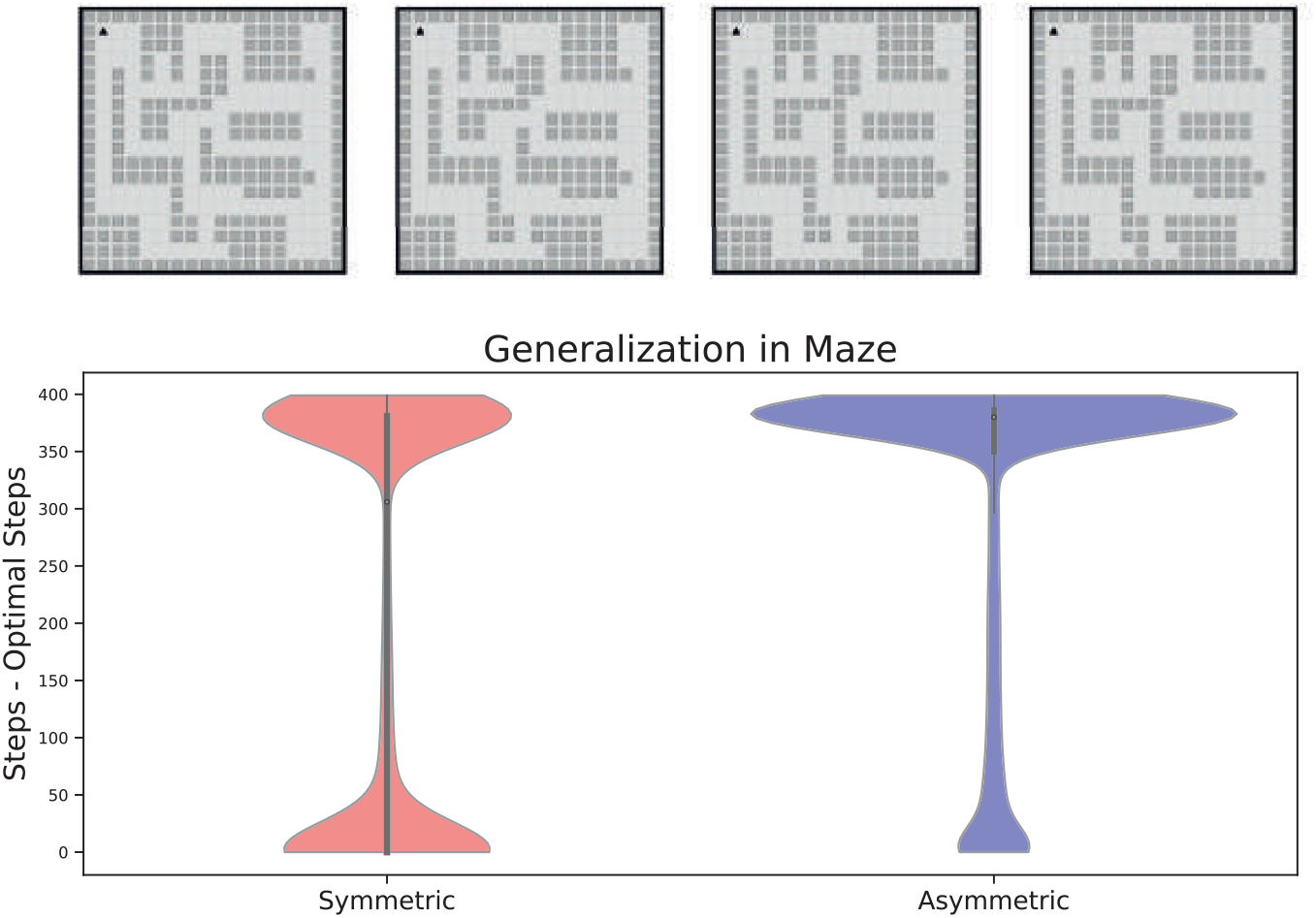
Generalization performance in maze task with blocked paths. **Top**: Grid world mazes used for generalization task. Leftmost maze was used for training, the other three environments for testing the generalization. **Bottom**: Violin plots show distribution of suboptimality (steps - optimal number of steps) of the agents when using the successor representation trained on one target and tested on another one. Training and test targets are randomly drawn from all possible states in the respective environments. The distribution for the symmetric agent is broader around 0, which indicates optimal generalization, and less pronounced at 400, which was the maximum number of steps allowed.

For the experiments in the maze, we used the same parameters for the maximum number of steps and the performance criterion, but higher values for the learning rate and the discount factor: *γ* = 0.9, *lr* = 0.999. This is because for lower parameter values the agents needed too many repetitions to converge.

Since all the environments we tested in the above setting have a symmetric nature, that is, their underlying state space is an undirected graph, we repeated the same generalization experiment in a setting where the state space is a directed graph, and hence travel time between two nodes is not symmetric. We constructed a simple graph with 17 nodes, which is essentially a tree graph with the addition of a directed edge from the lowest level to the highest. On this graph, we performed the same kind of navigation experiment. We used the same values for the learning rate and *γ* (0.1, 0.7), but reduced the number of episodes to 50 since the state space is smaller and hence can be learned faster.

#### 5.9.2 Policy Entropy

In Figure 5 we computed the entropy of the current policy of the agents after each episode of learning. As explained previously, we computed the current policy by applying the softmax-function to the q-values

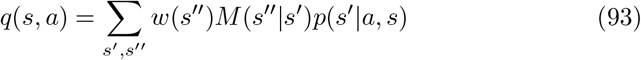

where *w* is the estimate of the reward vector and *M* is the current estimate of the successor representation. Then the policy is: *π*(*a* | *s*) = softmax(*q*(*s, a*)), and we computed the entropy of the policy averaged over all states as

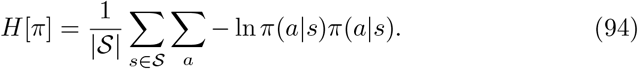

## 5.10 Acknowledgements

We thank William de Cothi, Tom George and Nikola Milosevic for helpful discussions.

## 5.12 Code Availability

All code and data used for this project is available at https://github.com/jakeck1/sympredlearning.

## A Learning Rule

In this appendix we are going to provide some additional insights into the learning rule we have employed. Furthermore, we will provide some informal derivation on the expected weights that should be learned under this learning rule. Recall hence our learning rule, which with parameters *α* = 1, *β* = 0 entails the update of synaptic weights *W* given pre- and postsynaptic activities *p*_*pre*_, *p*_*post*_ as

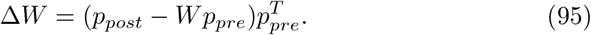

This learning rule can be seen as performing a sort of conditional expectation objective: At convergence, the expected update in synaptic weights, given the current activity, should be zero

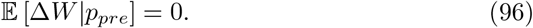

We have that this holds in general only if

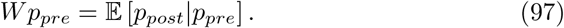

Note that *W* will also appear on the right hand side, as *p*_*post*_ will depend on *W* - thus, this equation still has to be solved for these weights. Now in some sense, this learning rule is very similar in spirit to classical TD-learning, since also there one tries to fulfill a conditional expectation equation (the Bellman equation) and works towards a solution by bootstrapping (Sutton and Barto, 2018). Indeed, also in Fang et al. (2023) this learning rule was derived starting from the assumption of TD-learning with function approximation.

Of course, the equation Equation 36 above will typically not have an exact solution, since conditioning on an input is in general a nonlinear operation. However, in special cases there will be a solution: If for example we have a conditional Gaussian relationship, that is *p*_*post*_ | *p*_*pre*_ ∼ 𝒩 (*p*_*post*_; *Ap*_*pre*_, *σ*), then an exact solution is possible: Then 𝔼 [*p*_*post*_ | *p*_*pre*_] = *Ap*_*pre*_ and hence *W* = *A* solves the equation - this example serves only as an illustration of possible solutions and there is no biological meaning intended. In general, one can understand the learning rule as enforcing a simple form of a predictive coding objective, where the weights are adapted in such a way that postsynaptic activity becomes predictable from presynaptic activity Huang and Rao (2011). Indeed, recall that the conditional expectation 𝔼 [*p*_*post*_|*p*_*pre*_] is the best prediction of *p*_*post*_, given *p*_*pre*_. Thus, if Equation 36 is satisfied, *W* indeed realizes this optimal prediction.

Let us now understand how the predictive coding rule learns successor representations.

### A.1 Learned representations through the predictive coding rule

Consider first a feedforward network with activities

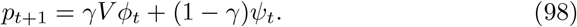

Here, *ϕ*_*t*_(*s*_*t*_) is the input through the feedforward weights *V* and *ψ*_*t*_(*s*_*t*_) is external input, both depending on the state *s*_*t*_ of a Markov process.

#### Lemma 1

*Applying the rule for the stationary weights from Equation 36 to the synaptic weights V and pre- and postsynaptic activities ϕ*_*t*_, *p*_*t*_ *respectively, yields the equation*

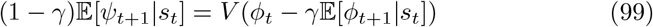

*Proof*. In this situation, we have *p*_*post*_ = *γV ϕ*_*t*+1_ +(1 − *γ*)*ψ*_*t*+1_, *p*_*pre*_ = *ϕ*_*t*_. Thus, the equation that the weights have to fulfill at convergence reads

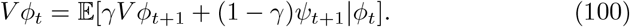

Now as we assume an injective relationship of states *s*_*t*_ and features, we may replace the conditioning on presynaptic activity by conditioning on hidden states *s*_*t*_. Then pulling the weights *V* out of the expectation due to linearity and then brining all terms involving *V* to one side yields the result.

We assume the case of a finite state space, where we can write 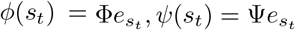, for matrices Φ, Ψ which we assume invertible. The equation is then

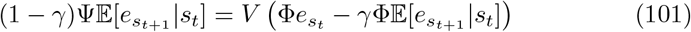

It is then not hard to see that a solution is

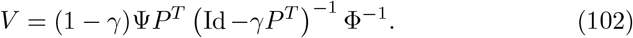

But this is now just successor features (up to one timestep) in another basis: We have that applying *V* to a feature vector *ϕ*_*t*_ yields

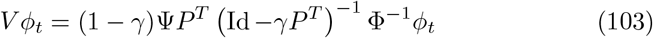

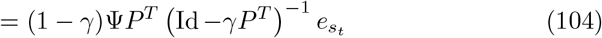

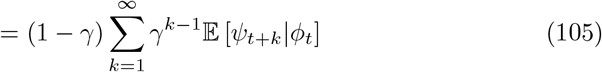

which is hence indeed the successor representation of *ψ* conditional on the features in *ϕ*. In particular, plugging this solution for *V* into equation Equation 98 then just includes the current timestep into the sum

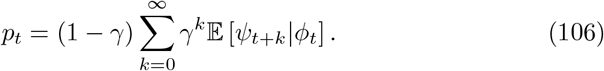

The situation for the recurrent net is maybe even simpler (cf. Fang et al. (2023)): Assume a network with activity

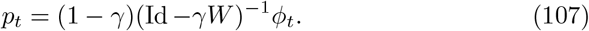

Then Equation 36 for the weights *W* becomes

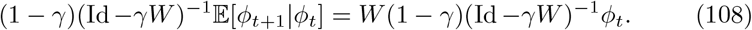

Since *W* and (Id − *γW*)^−1^ commute, it is again not hard to see by similar algebraic manipulations as before, that a solution is

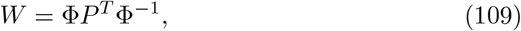

and hence when plugging *W* back into the equation for *p*,

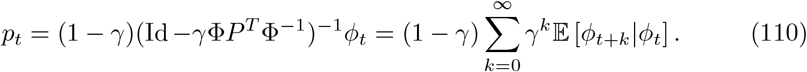

Thus we see that now *W* encodes a successor representation for features *ϕ*, conditioned on *ϕ* itself.

Finally, we want to ask what happens when the input to the feedforward-layer itself is also a successor representation, under the same prediction horizon *γ*. That is, when the matrix of input features is 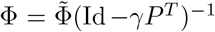, for some other set of features 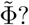 In this case the matrix *V* takes the particularly easy form 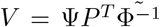, which is just the one step prediction of features Ψ from features 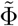 (analogous to the recurrent case above). We see that still in this case *p*_*t*_ will encode successor features, just conditional on the basis 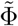: Plugging in the solution for *V* into the equation for *p*_*t*_ yields

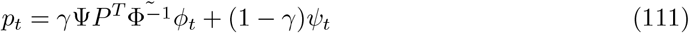

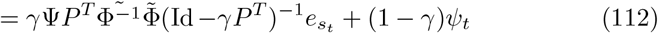

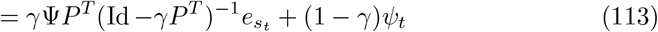

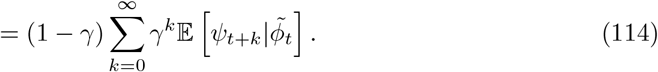

This shows that when one has a feedforward network which itself receives successor features as inputs, the result will again be successor features.

The limits derived in this section give the right intuition and are indeed the correct ones, we give a more formal analysis of the convergence below.

## B Backward shifts of successor features

Here, as mentioned in subsection 5.5, we want to prove backward shifting of successor features which are Gaussian, but let us first make some general remarks. Recall that we are assuming an idealization of a linear track as the real line ℝ, where an agent is moving on. We are considering a feature *ϕ* : ℝ → ℝ and want to study the shift of it’s center of mass, defined as the normalized mean 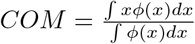

Since the feature in our case is obtained as inputs from other neurons, it makes sense to assume that *ϕ* is non-negative. Without loss of generality, we also assume that the feature is normalized, that is *ϕ* is a probability density, and the center of mass simply becomes ∫*xϕ*(*x*)*dx* = 𝔼_*ϕ*_[*x*]. Furthermore, when we normalize the successor operator (which is simply the Laplace transform) by multiplying by *γ*, that is by letting

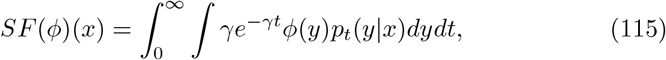

then we have that *SF* (*ϕ*)(*x*) = 𝔼_*q*(*y*|*x*)_[*ϕ*(*y*)], with 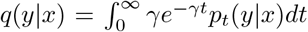 the density obtained from transforming *p*_*t*_(*y* | *x*) (one can think of this as the transition density from *x* to *y* when time *t* is sampled from an exponential distribution). In general, it might be difficult to study the centre of mass, because for example *SF* is not necessarily normalized anymore.

However, the case we consider is more benign, so let us turn to the special case now after these general remarks. Assume that we have Gaussian features, that is

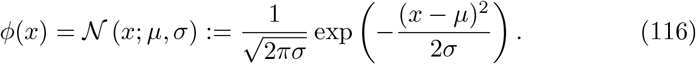

Hence, the centre of mass is simply the mean *µ*. Let the dynamics be a Brownian motion with a constant drift *v*, that is

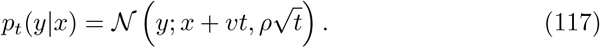

In particular, as a function of *x* we can write this as

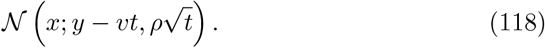

Then integrating the transition density and *ϕ*, just yields the marginal distribution, which by the well known properties of Gaussians is again a Gaussian

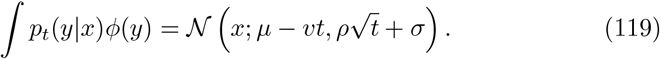

Thus, the successor feature is then given by

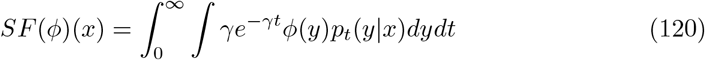

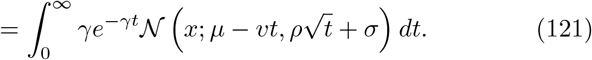

In particular, the centre of mass is

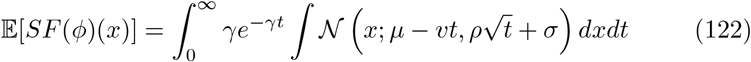

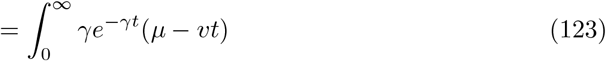

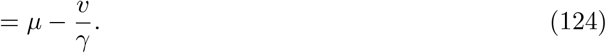

Thus, we observe again, just as in the example considered in the main text, a shift which depends on the predictive time scale factor *γ* and the velocity *v* - with a bigger *γ* or a smaller *v* leading to less shift. In particular, if the velocity is zero, then no shift should be observed.

## C Eigenvectors of the Successor Representation

The eigenvectors of the successor representation matrix (Id − *γP*)^−1^ are of special importance in the theory both in computational neuroscience as well as in reinforcement learning. In the latter, these eigenvectors are used to construct a natural basis for functions on the state space, which yield a representation of the large-scale geometry of the state space and can be used for example for option discovery (Machado et al., 2017b,a; Mahadevan and Maggioni, 2007). In neuroscience in turn, these eigenvectors have been used as a model of grid cells, since in spatial settings there are remarkable similarities of some of the eigenvectors to grid cells observed in the entorhinal cortex (Stachenfeld et al., 2014, 2017). The key insight to why these eigenvectors are particularly useful to represent the state space is that they are essentially the eigenvectors of a Laplace operator (Stachenfeld et al., 2014). On an undirected graph with adjacency matrix *A*, the random-walk Laplacian is defined as

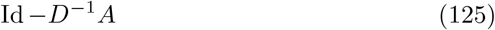

where *D* is the diagonal matrix with entries 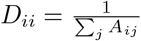. There are other ways to define a Laplacian (Chung, 1996), but on an undirected graph they are all related (Jost and Mulas, 2019). In Equation 125, the matrix *P* := *D*^−1^*A* corresponds to the transition probability matrix of arandom walk where the neighbors of a vertex are visited with the same probability. It is then not hard to see that the random walk Laplacian and the successor representation under *P* share the same eigenvectors (although for different eigenvalues). These eigenvectors and associated eigenvalues have many desirable properties: they are invariant under certain symmetries (automorphisms) of the graph (Mahadevan and Maggioni, 2007), they can be used to cluster the graph (Von Luxburg, 2007) and they represent slowly varying/smooth features (Sprekeler, 2011). Many of the technical results on graph Laplacians hinge on self-adjointness (i.e., symmetry), which is why they do not directly translate to the setting of directed graphs. In this situation, it is thus common to include some form of symmetrization - in fact, from the beginning in RL it was proposed to use Laplacian eigenfunctions under a symmetrization (Mahadevan and Maggioni, 2007). There are at least two straightforward ways how to do this: either, one can treat each directed edge as an undirected edge - this corresponds to constructing a new adjacency matrix 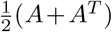, and was for example used in Mahadevan and Maggioni (2007) to obtain basis functions for RL. The other possibility is to use the information from the stationary distribution, and symmetrize as 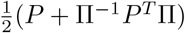 this leads to the so called ‘Chung-Laplacian’ (Chung, 2005); also this Laplacian construction has been successfully applied in RL (Johns and Mahadevan, 2007; Wu et al., 2018). We have seen that our symmetrized learning rule in the stationary setting will learn the successor representation under *P* +Π^−1^*P*^*T*^ Π, and therefore eigen-values constructed from it would correspond to those of the Chung-Laplacian. However, in practice stationarity might not be a valid assumption - this is for example the case in our simulations, where episodes are terminated on reward. In this case, the symmetric representation that is learned is actually more closely related to the former case.

## D Convergence proofs

In the following, we will give convergence proofs for both the neural network model and the TD-learning setting. In both situations, we will use the same strategy as in Dayan and Sejnowski (1994). Therein, they use a classical result from stochastic approximation theory (Kushner and Clark, 2012). We state now a formulation of this result close to the one used in Dayan and Sejnowski (1994), but note that this omits some detail.

### Theorem 1.

*Let Z*_*n*_ ∈ ℝ^*n*^ *be a sequence of random variables, which fulfill*

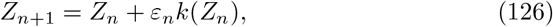

*where k is a stochastic function of Z mapping to* ℝ^*n*^, *and ε*_*n*_ ∈ (0, 1) *is a sequence such that* 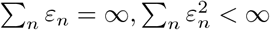. *Define* 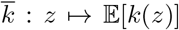, *assume that Z*_*n*_ *is bounded almost surely and that* Var 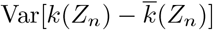 *is bounded. Let A*^0^ *be the set of asymptotically stable equillibria of the differential equation*

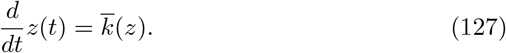

*Then if A*^0^ *is nonempty, Z*_*n*_ → *A*^0^.

As in Dayan and Sejnowski (1994), we will not verify the boundedness conditions, because these could be enforced by projecting the sequence back into a bounded region if necessary i.e., one could modify the update equations to include a projection step that enforces staying in a certain bounded region. Then, all quantities above are bounded and hence this new sequence converges to *A*^0^ if *A*^0^ is in that region.

### D.1 Convergence of Model

In the following, to avoid using too many superscripts, we change notation compared to the main text, and set *W* = *W* ^1^ the recurrent weight,*V* = *W* ^2^ the feedforward weight, *p* = *p*^1^, *q* = *p*^2^ the population activities, *ϕ* = *ϕ*^1^, *ψ* = *ϕ*^2^ the inputs. Furthermore, we use the notation

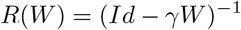

and recall our definition

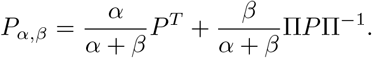

Furthermore, recall that we assume our neural activities *p, q*, given inputs and synaptic weights are

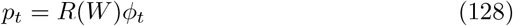

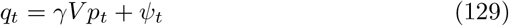

Consider the update rule for the system

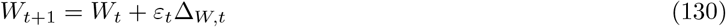

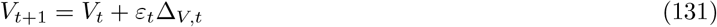

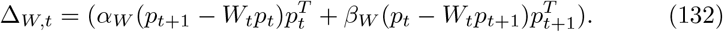

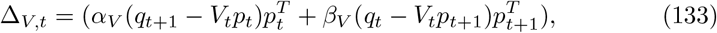

As stated in the theorem above, we can show convergence to *W* ^*^, *V* ^*^ if we can show that the differential equation

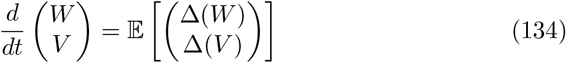

has an asymptotically stable fixed point at *W* ^*^, *V* ^*^ - to do so, we will study eigenvalues of the Jacobian at putative fixed points. Recall that for an autonomous dynamical system *dx/dt* = *F* (*x*) and some fixed point *x*_0_, if the linearized version *du/dt* = *Df* (*x*_0_)(*u*) of the system is asymptotically stable at *x*_0_, so is the original system. In turn, if the Jacobian *DF* (*x*_0_) of the system at that fixed point has eigenvalues with all negative real parts, the linearized system is asymptotically stable. This lays down our strategy to prove stability.

First, we compute the right hand side of the differential equation Equation 134. We have approximately (assuming for the expectation that *W*_*t*+1_ ≈ *W*_*t*_, *V*_*t*+1_ ≈ *V*_*t*_):

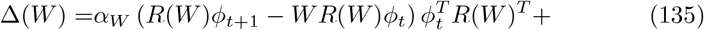

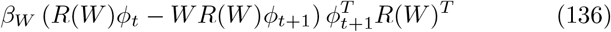

and (recall *q* = *γ*_*V*_ *V p* + *ψ*)

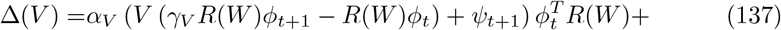

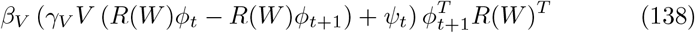

We now assume that the inputs *ψ*_*t*_, *ϕ*_*t*_ are in stationary distribution (we state below a condition under which this holds) and then take the expectation to obtain

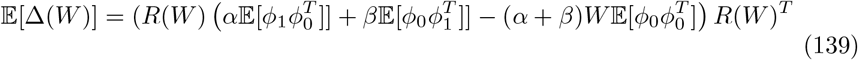

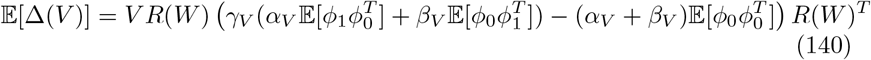

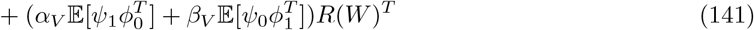

where we have used that *W* and *R*(*W*) commute. We now will find the putative fixed points oft these equations by finding their zeros. For the first equation, setting the innermost term zero, we see that this equation has an equilibrium at *W* ^*^ if

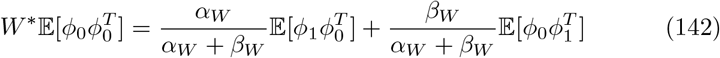

We see that this equation essentially prescribes *W* to implement a weighted average of the linear regression of *ϕ*_0_ against *ϕ*_1_ and vice versa. Indeed, if 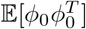 is invertible one sees that it becomes

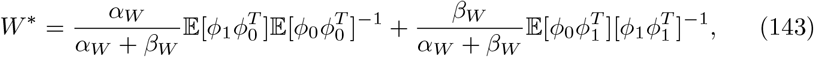

and 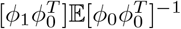 is for example the regression weight for linearly predicting *ϕ*_1_ from *ϕ*_0_.

Setting now the second equation to zero yields the relation which has to hold at a putative fixed point *V* *:

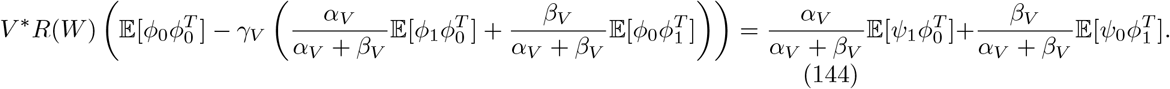

We now want perform a stability analysis of the equations at an equillibrium (*W* ^*^, *V* ^*^). Since the dynamics of *W* do not depend on *V*, we can look at the respective equations separately and if both are stable, the whole system is stable. The easier part is the equation for *V* : It is affine linear in *V*, that is we only have to study the eigenvalues of the matrix

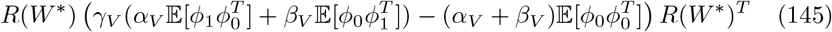

In fact, using lemma 3, we only have to study the eigenvalues of the inner term 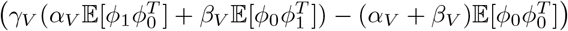. According to lemma 6, these have all negative real parts, hence we have stability for this equation at any equilibrium.

The stability of the differential equation for *W* is slightly more delicate. Let *F* denote the right hand side of the differential equation for *W*. We want to find its Jacobian. First, we want to calculate the derivative of *F* (in the Frechet sense) at *W* ^*^, as a linear map applied to a matrix *U*. We have that

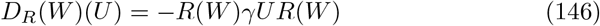

and thus by the chain rule

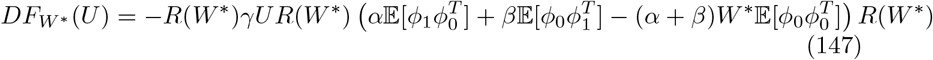

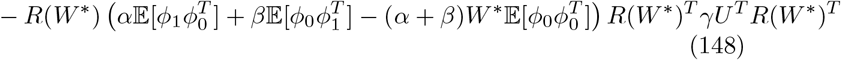

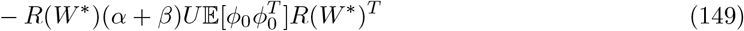

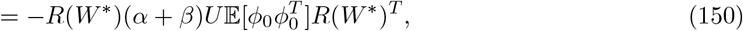

where the first two terms vanished at *W* ^*^ because by assumption

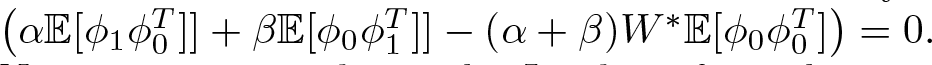

Now, we want to obtain the Jacobian from the expression above, that is we want to find a matrix that represents this derivative. To achieve this, we may vectorize the matrix *U* using the vectorization operator vec, which transforms a matrix into a vector by stacking the columns on top of each other. Using the well known identity

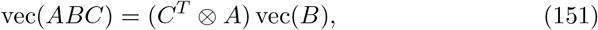

where ⊗ denotes the Kronecker product of matrices, we thus have

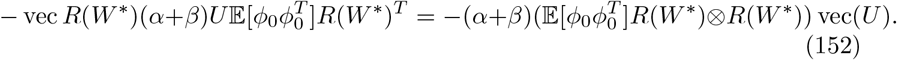

This shows that the Jacobian is given by the matrix

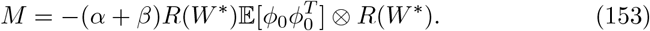

Now, for the Kronecker-product of two matrices, the eigenvalues are given as the products of the eigenvalues of the individual matrices. That is, if *λ*_*i*_ are the eigenvalues of 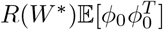 and *µ*_*j*_ are the eigenvalues of *R*(*W* ^*^), the eigenvalues of *M* are given by

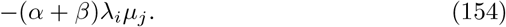

Now there seems to be in general *no* guarantee that these all have negative real parts and thus the matrix *M* be stable for all choices of *α, β, ϕ*.

However, we can obtain a results in the symmetric case, under some assumptions (which will hold in the case we are interested in):

#### Proposition 1.

*For* 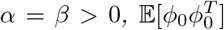 *invertible, and the real part of any eigenvalue µ of W* ^*^ *fulfilling* Re(*µ*) ≤ 1 *the matrix M is stable, that is all eigenvalues have negative real parts*.

*Proof*. First, we want to consider the eigenvalues of 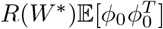. We note that by the defining relation of *W* ^*^ as a fixed point we obtain that

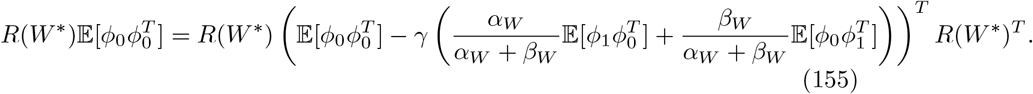

According to lemma 6 and 3, the eigenvalues of the right hand side have positive real parts, and for *α* = *β*, the matrix is symmetric hence the eigenvalues are positive reals. On the other hand, the eigenvalues of the matrix *R*(*W*)^*^ = (Id − *γW* ^*^)^−1^ have positive real parts by assumption and lemma 4. Thus, also the products of the eigenvalues of the two matrices have positive real parts, which proves the claim.

We have thus confirmed that in the symmetric case, it is possible to have stability (and thus convergence) at some (*W* ^*^, *V* ^*^). Below, we will now assume a specific structure of the inputs which will link to successor features and which will make it possible to state the existence of a solution. We will now assume

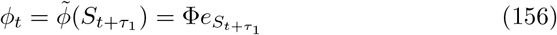

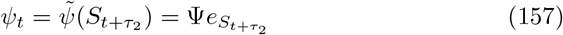

where *S*_*t*_ is the state process, which we assume in stationary distribution, 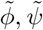 are injective functions, the matrices Ψ, Φ have the respective values of these functions as their columns, The *τ*_*i*_ indicate that the two inputs could have a temporal offset, we will however set these to zero in the sequel. We note at these points that these choices are made to relate to the definition successor features, and other choices of input could equally yield a similar result.

With the above definitions, we have

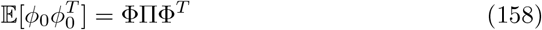

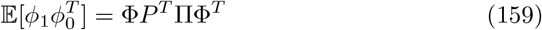

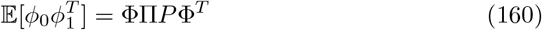

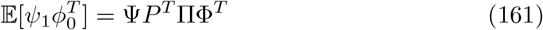

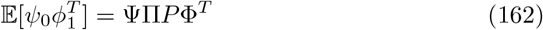

where *P* is the transition matrix of *S*_*t*_ and Π is again the diagonal matrix of the stationary distribution. Plugging these expressions into the relation we derived for *W* ^*^ yields

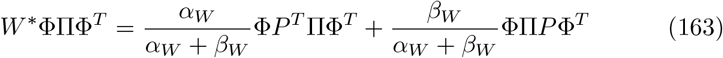

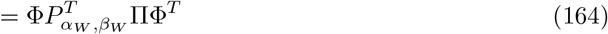

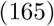

where it is now easy to see that all possible solutions for *W* ^*^ are of the form

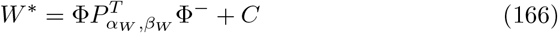

where Φ^−^ denotes the pseudoinverse, and *C* is a matrix with *C*Ψ = 0.

Similarly, for *V* ^*^ we obtain the relation

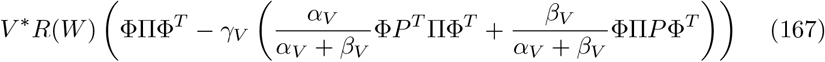

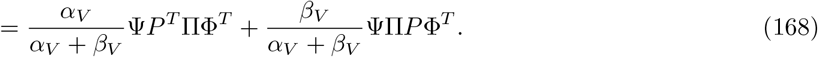

which may be written as

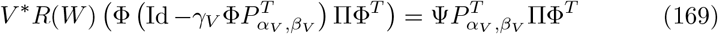

It is then again clear that all possible solutions for *V* ^*^ are of the form

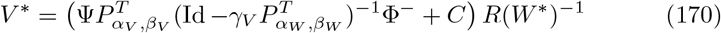

where *C* is again an arbitrary matrix having Φ in its kernel.

We note that with this particular form of *W* ^*^, we can indeed achieve stable solutions: 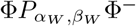 has eigenvalues either being zero or corresponding to eigenvalues of 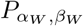, which is a stochastic matrix and hence has eigenvalues in the unit circle, such that as long as the matrix *C* has eigenvalues with real part lesser or equal than 1, the requirements of our proposition above are met and we have a stable equilibrium. In the particular special case where Φ is invertible, we get unique stable solutions.

#### D.1.1 Limits of activities

With these particular solutions, we can understand the form the activities of the populations will take. We will plug in the limits of the weight matrices into the equations

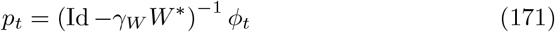

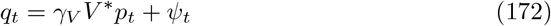

First, we recall that we assume inputs of the form *ϕ*_*t*_ = *ϕ*(*S*_*t*_). Second, we note that the action of the matrices on the space spanned by Φ is the same for all solutions *W* ^*^, regardless of the additional matrix *C*: As *C*Φ = 0, whenever our input is in the space spanned by the features, we have 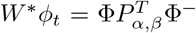. Now, since Φ^−^ acts as an inverse on this space, we have that

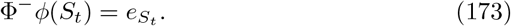

Hence, in particular

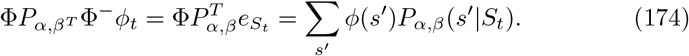

Thus, we have shown that

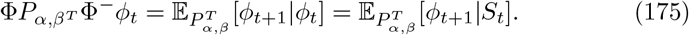

The same arguments holds for any powers of the matrix *W* ^*^: Indeed, taking

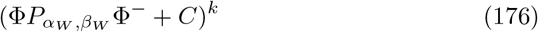

will result in a sum of mixed terms of powers of the two matrices in the sum. However, whenever *C* appears to the left of Φ, the matrix will be zero, and whenever *C* is the rightmost term, the matrix again has the span of the columns of Φ in its kernel. Thus, one may consider only the terms

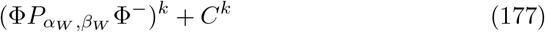

and then perform exactly the same calculation as above. This then yields

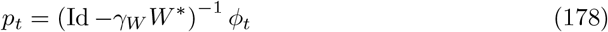

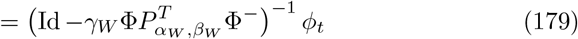

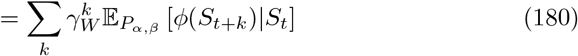

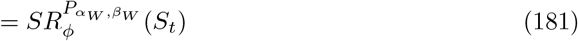

which shows that indeed *p*_*t*_ encodes the successor representation of the function *ϕ*.

Now for *q*_*t*_,first recall that for *V* ^*^ we obtained the limit

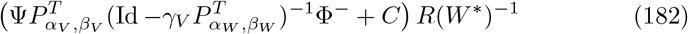

where *C* is a matrix with the span of Φ in its kernel and *R*(*W* ^*^) = (Id −*γ*_*W*_ *W* ^*^)^−1^. Now plugging this in into the equation for *q*_*t*_ yields

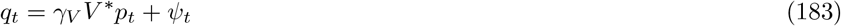

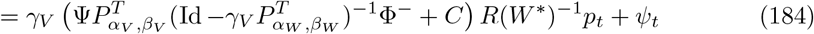

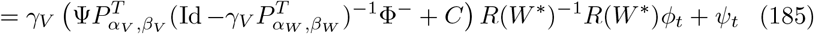

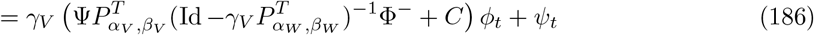

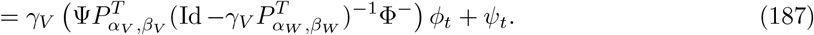

Then, note that

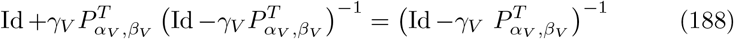

as can be seen immediately by multiplying both sides by 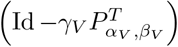.

Further note that 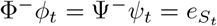 by definition. Using this, we have

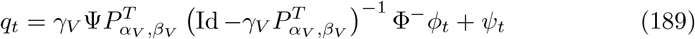

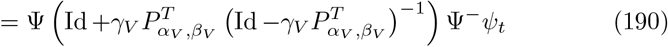

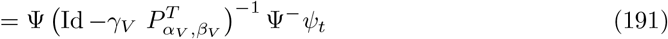

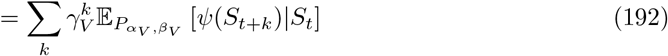

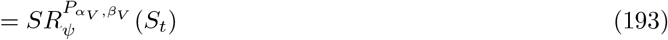

which shows that also *q* encodes a succesor representation.

## E Convergence of TD learning

Here we show in a very similar manner that classical TD-learning with the inclusion of parameters *α, β* learns the successor representation under the modified probabilities. In fact, one can see classical TD-learning as a one layer feedforward network implementing the learning rule/

Consider the following modified TD-learning rule for the parameter *θ*_*t*_ parametrizing 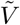

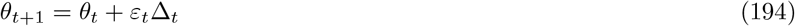

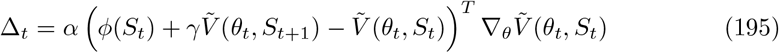

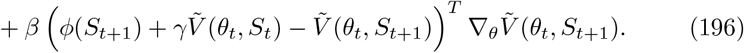

We can see this as a semi-gradient descent with respect to the loss

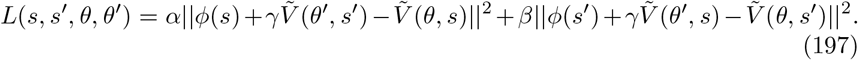

Now assume the linear and tabular case, that is

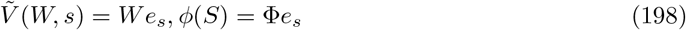

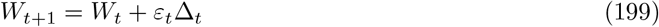

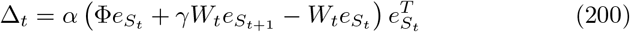

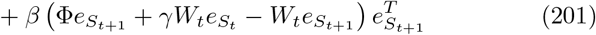

Note that we can also write the updates as

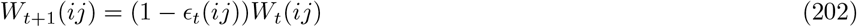

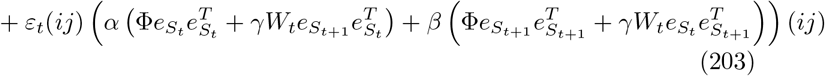

where *ϵ*_*t*_(*ij*) = *ε*_*t*_(*α*𝕀[*S*_*t*_ = *j*] + *β*𝕀[*S*_*t*+1_ = *j*]).

We have

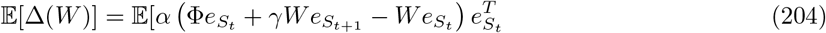

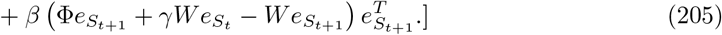

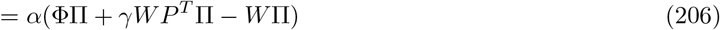

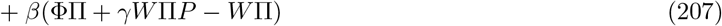

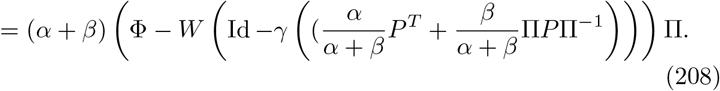

Setting to zero, we obtain

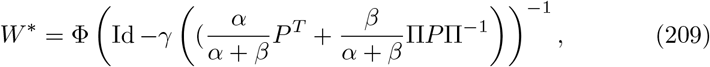

which is precisely the successor representation under *P*_*α,β*_. The stability of the differential equation, which is linear, is determined by the eigenvalues of the matrix

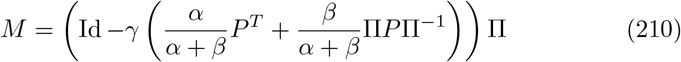

Using again lemma 6 below this has all positive eigenvalues, which proves the stability and hence the convergence.

## F Auxiliary results

Here we state some lemmas we used for the convergence proofs above, these should all be standard results, but we prove them here for convenience.

### Lemma 2.

*Let M* ∈ ℝ^*n×n*^ *(not necessarily symmetric). Then if M is positive definite in the sense that for any* 0 *have positive real part. x* ∈ ℝ^*n*^, *x*^*T*^ *Mx >* 0, *all eigenvalues of M*

*Proof*. Let *λ* ∈ ℂ be an eigenvalue of *M* with eigenvector *z* ∈ ℂ^*n*^. Then on the one hand we have

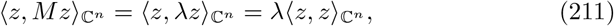

and on the other hand

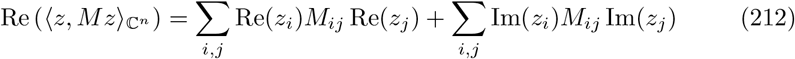

Now since 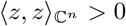, if *M* is positive definite, the right hand side of the last equation is positive and this implies that *Re*(*λ*) *>* 0.

### Lemma 3.

*Let A, B be in* ℝ^*n×n*^ *and B be invertible. Then A is positive definite in the sense of the previous lemma if and only if so is BAB*^*T*^.

*Proof*. Since *B* is invertible,

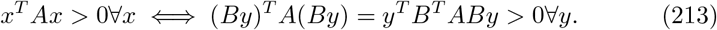

### Lemma 4.

*Let Q be a matrix Re*(*λ*) *<* 1 *for all its eigenvalues λ* ∈ ℂ. *Then Id* − *Q has eigenvalues with all positive real parts, is invertible, and the inverse also has eigenvalues with all positive real parts*.

*Proof*. The eigenvalues of *Id* − *Q* are of the form 1 − *λ*. Hence we have Re(1 − *λ*) *>* 0. This then implies that *Id* − *Q* is invertible. The last part follows since inversion maps the positive half-plane to the positive half-plane.

### Lemma 5.

*Let A be a square matrix and A*^*^ *be it’s adjoint with respect to some inner product* ⟨·, ·⟩. *Then we have, denoting by σ the spectrum*,

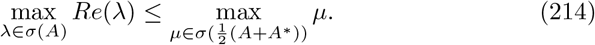

*Proof*. We note that for an eigenpair (*v, λ*) of *A*

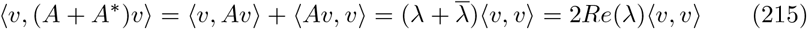

Hence, by the Courant-principle

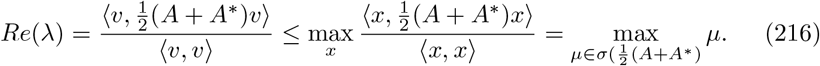

### Corollary 1.

*Let t* ∈ ℝ, *P a stochastic matrix with stationary distribution π, and* Π = diag(*π*). *The matrix*

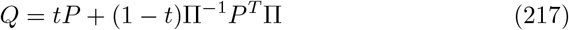

*has Re*(*λ*) ≤ 1 *for all its eigenvalues λ* ∈ ℂ.

*Proof*. Consider the inner product ⟨*x, x*⟩ _Π_ = *x*^*T*^ Π*x*. With respect to this inner product, the adjoint of *Q* is

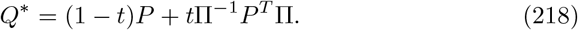

Hence,

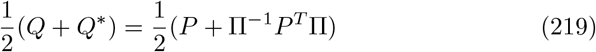

which is a stochastic matrix and therefore has largest eigenvalue 1.

### Lemma 6.

*Let ϕ*_0_, *ϕ*_1_ *be random vectors distributed equally. Let furthermore t* ∈ ℝ *and γ* ∈ [0, 1). *Then the matrix*

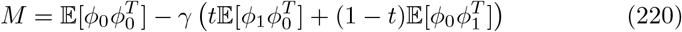

*has eigenvalues with all nonnegative real parts. If furthermore* 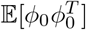 *has full rank, then the real parts are all positive*.

*Proof*. Let *x* be any nonzero vector. Note that we have

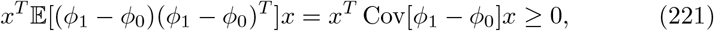

hence

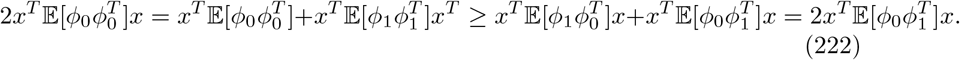

From this follows *x*^*T*^ *Mx* ≥ 0 with equality only if *x* is in the kernel of 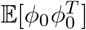 and hence the claim.

## G Symmetrized distribution is closer to uniform distribution

It is intuitive, that a reversible distribution of transitions is closer to the uniform distribution. The Kullback-Leibler divergence yields a particularly easy way to prove this intuition. Assume our state space S is a undirected graph *G* = (S, *E*), that is, we are in a deterministic setting, and if one can go from *s*_1_ to *s*_2_ then one can also go back. Then, together with the stationary distribution *π*(*s*), the transition probabilities *p*(*s* | *s*^*′*^) define a joint probability on the set of edges (with orientation) of the graph, i.e.

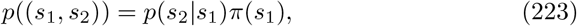

with (*s*_1_, *s*_2_) denoting the directed edge from *s*_1_ to *s*_2_. That is, if we take the transition probabilities

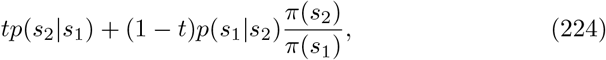

then the joint distribution would be simply

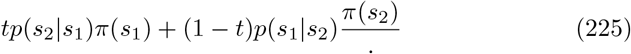

Now, if we were to observe transitions uniformly, then we have

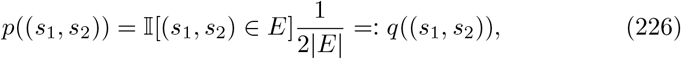

where the factor 2 appears because we consider each undirected edge twice. Note that this is the joint probability that is obtained when one classically defines the random walk on an undirected graph with

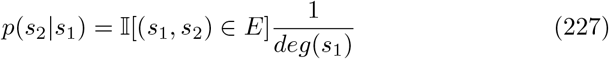

with *deg*(*s*_1_) = ∑_*s*_^*′*^ 𝕀[(*s*_1_, *s*^*′*^) ∈ *E*] the degree. Indeed, weighting the above equation with the stationary distribution 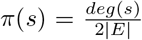 yields the uniform distribution over edges.

Now, the Kullback-Leibler divergence between any probability distribution and the uniform distribution is just

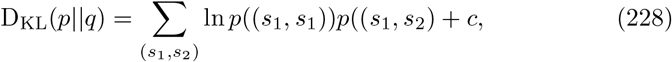

with *c* not depending on *c* -i.e, it is just given by the negative entropy of *p*. Hence, the following lemma shows that indeed choosing 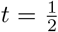 in Equation 224 leads us closest to uniform.

### Lemma 7.

*Let X* ∈ 𝒳 *be a random variable with density p, where* 𝒳 *admits an involution, that is a bijective, measure-preserving map σ* : 𝒳 → 𝒳 *such that σ*^2^ = *Id. Let H*(·) *be the entropy. Then for any* 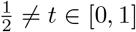:

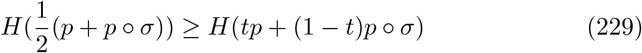

*and equality holds iff both densities are equal*.

*Proof*. By the strict concavity of entropy and the convexity of the set {*tp* +(1− *t*)*p* ◦ *σ*|*t* ∈ [0, 1]}, there is a unique maximum. Differentiating with respect to *t* yields

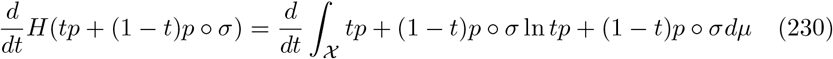

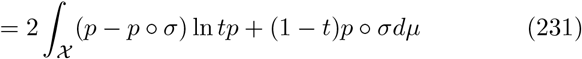

Now at 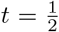, the term ln 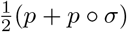 is invariant under *σ*, while the term *p* − *p* ◦ *σ* is skew-symmetric under *σ*. Hence, the integral vanishes.

### G.1 Proof that symmetrization of optimal policy is stable

Here, we want to show that an optimal policy remains optimal even if the value function is computed under symmetrized transition probabilities. Before we do so, we will however first state an useful identity for the successor representations.

#### Definition 1.

*Under a fixed policy π, for S*_*t*_ *a random walk with transition probabilities P* (*s*^*′*^|*s*) = *p*(*s*^*′*^|*a, s*)*π*(*a*|*s*), *we define the first hitting times*

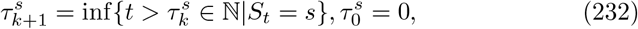

*where if the case that the set is empty we set the value to* ∞. *For a given pair of states s, s*^*′*^ *we define*

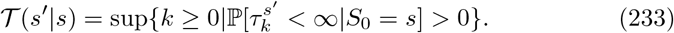

*Note that* 𝒯 (*s*^*′*^|*s*) *only takes the values* {0, 1, ∞}.

It is easy to see that by, the Markovian property, the distribution of these stopping times does not change, in the sense that

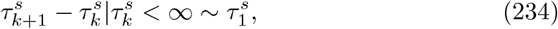

where for the left hand side we used the convention ∞ − *c* = ∞ for any finite *c*. Using this, we have the following expression for the Successor representation in terms of stopping times:

#### Proposition 2.

*For the successor representation M* = (*Id* − *γP*)^−1^ *we have*

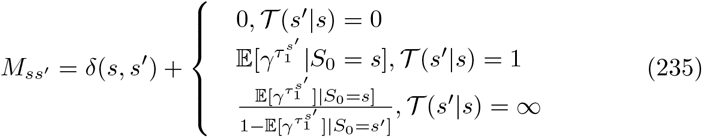

*Proof*. We note that

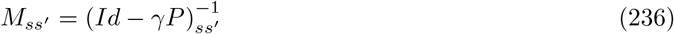

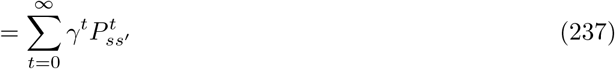

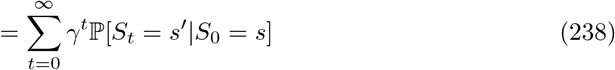

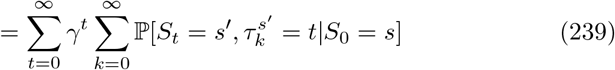

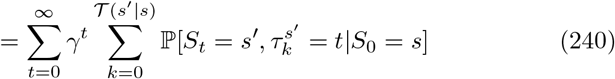

where we have simply included a sum over all possible values for the hitting times and then included only the cases where those are finite.

Now note that in the above sum

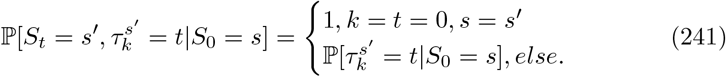

Hence we get

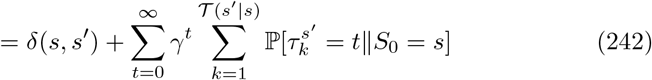

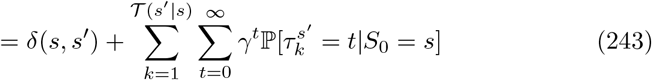

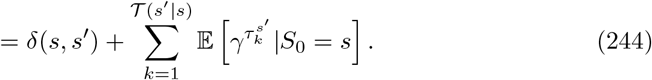

In particular, if 𝒯 (*s*^*′*^|*s*) = 0, we have *M* (*s*^*′*^|*s*) = *δ*(*s, s*^*′*^). In the case that 𝒯 (*s*^*′*^|*s*) = 1, we get

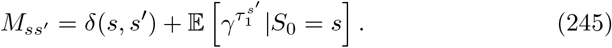

Now, for the remaining case 𝒯 (*s*^*′*^|*s*) = ∞ first note that

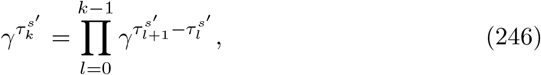

and that for *l >* 0 we have 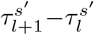 is independent of *S*_0_ and since at 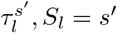, we have

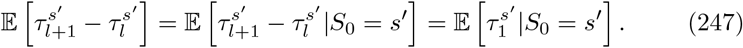

Hence,

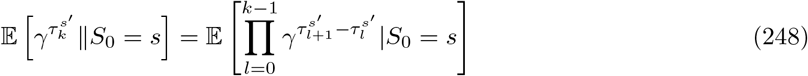

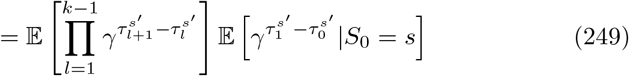

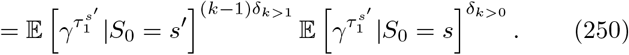

Plugging this into the sum formula yields

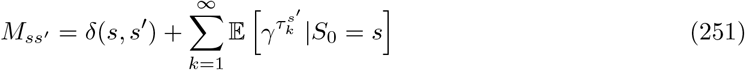

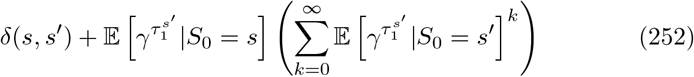

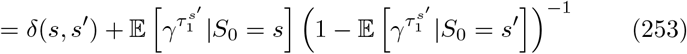

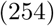

as claimed.

#### Corollary 2.

*In the setting with a unit reward e*_*s*\*_, *the value function is completely determined by the expected γ to the power of the first hitting time, that is*

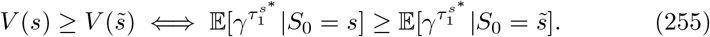

In particular, consider the deterministic setting - that is, the transitions *p*(*s*^*′*^| *a, s*) are deterministic and hence the state-space becomes a directed graph, where a directed edge exists from state *s* to state *s*^*′*^ if there is an action *a* which (deterministically) leads from states *s* to state *s*^*′*^. The task to navigate to state *s*^*′*^ is inherently finite/absorbing, but we can extend the time horizion to infinity by simply allowing an action that stays in the same state as one is (which hence is the optimal action once one is at the target state). In this deterministic setting, an optimal policy has to always decrease the distance to the target - hence, we have under an optimal policy, where the target is *s*^*^,

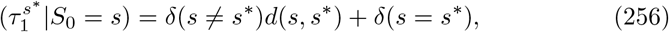

where *d*(*s, s*^*′*^) is the shortest number of steps necessary to get from *s* to *s*^*′*^ (which is not necessarily symmetric). In particular, every non-target state *s*^*′*^ should never be returned to, and starting from state *s*, only states lying on shortest paths from *s* to *s*^*^ should be visited (where some of those shortest paths may still not be visited by the policy, as there might be multiple such paths). In summary:

#### Corollary 3.

*In the directed graph setting navigating to target s*^*^, *for an optimal policy π*^*^, *the successor representation becomes*

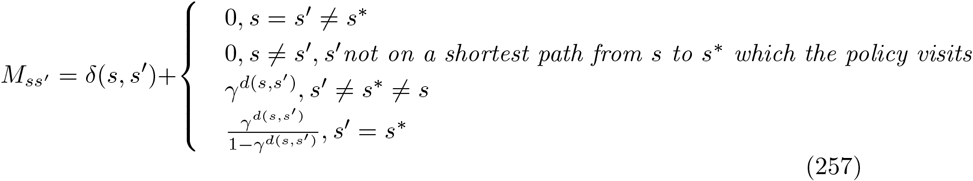

Having understood this simple structure of the *SR* under an optimal policy, we can also understand why a symmetrization does not impede solving the task. First, we have to slightly generalize our definition of the symmetrization of transition probabilities, as the one we used in our biological model only works in the case where the resulting Markov chain is ergodic. This is not the typical situation in navigation problems. Rather, in navigation tasks, under an optimal policy, one will navigate towards a goal on the shortest possible path and then typically the episode ends. Thus one could on the hand imagine an absorbing state at the target, but this then does not lend itself nicely to the generalization setting where the target should change but the environmental dynamics should stay the same. Instead, one could then not introduce an absorbing state in the environment transitions per se, and just include a self-transition at the target state - but still stop the episode accordingly when the reward is reached.

We thus would want to define the reverse process in this case as a process which always moves away from the target state *s*^*^, until it reaches a state with maximal distance to the target, where it stays (or rather, the episode ends) - hence mirroring the behaviour of the optimal policies. That is formally, if *P* are the transition probabilities under the optimal policy, and *d*_max_ = max_*s*_ *d*(*s, s*^*^)

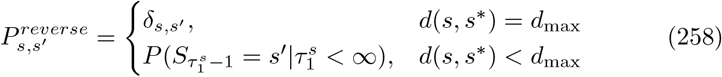

The symmetrized process is then the one with transition probabilities given by

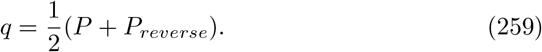

The value function induced by *q* for the navigational problem (i.e. reward vector *e*_*s*\*_) is then

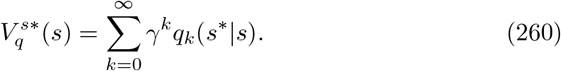

Recall that a policy *π* is optimal for a value function *V* if the policy always selects actions which maximize the value function. We have

#### Proposition 3.

*Let π be an optimal policy in the deterministic navigation problem with target state s*^*^ *and let q be the corresponding symmetrized process. Then π is also an optimal policy for* 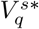.

*Proof*. First, let us define the sets

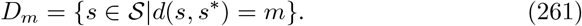

It is obvious that under the optimal policy, for the forward process, we have

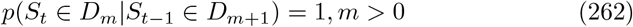

and

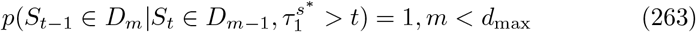

This means that for the symmetrized process, with transition probabilities 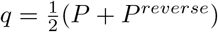, we actually have

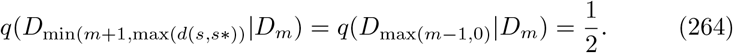

That is, the symmetrized process with probability 1*/*2 either increases or decreases the distance to the target state *s*^*^ by 1, with exception at the boundaries (i.e. at maximal distance or zero distance), where it stays with probability 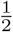. Now consider the value-function under *q*, which depends on the sum of the *k*−step probabilities *q*_*k*_. But the *k*− step probabilities can be written as

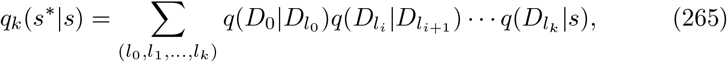

where the sum runs over all possible alignments such that one can reach *s*^*^ after *k* steps from *s*. This implies that the value-function can be completely determined by looking at the transition probabilities on the coarse-grained state space of level sets of the distance function, which corresponds to a line graph which self-loops at the ends. To be more precise, the value of *s* under *q* is determined simply by

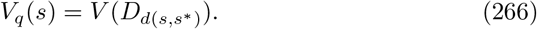

This tells us that the value of the state is a function only of the distance to the target state. Now we only need to show that this function decreases strictly with distance. To do this, we can simply study the line-graph, or rather the process with transition matrix

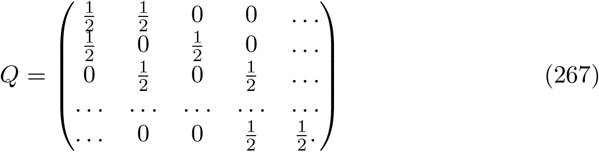

Now let *d*(*s, s*\*) *< d*(*s*^*′*^, *s*\*). It is then clear that 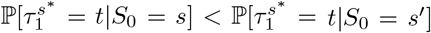, since any path from *s*^*′*^ to *s*^*^ has to go through *s*. Hence, recalling proposition 2, also

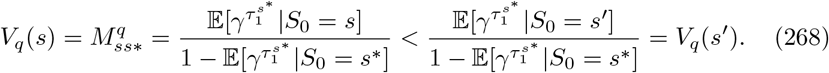

This shows that the indeed the value-function decreases strictly with distance, and hence the optimal policy *π* which we started with, also strictly decreases *V*_*q*_ and hence also is an optimal policy for *q*, which concludes the proof.

## H Supplementary Figures

**Figure H.1:**
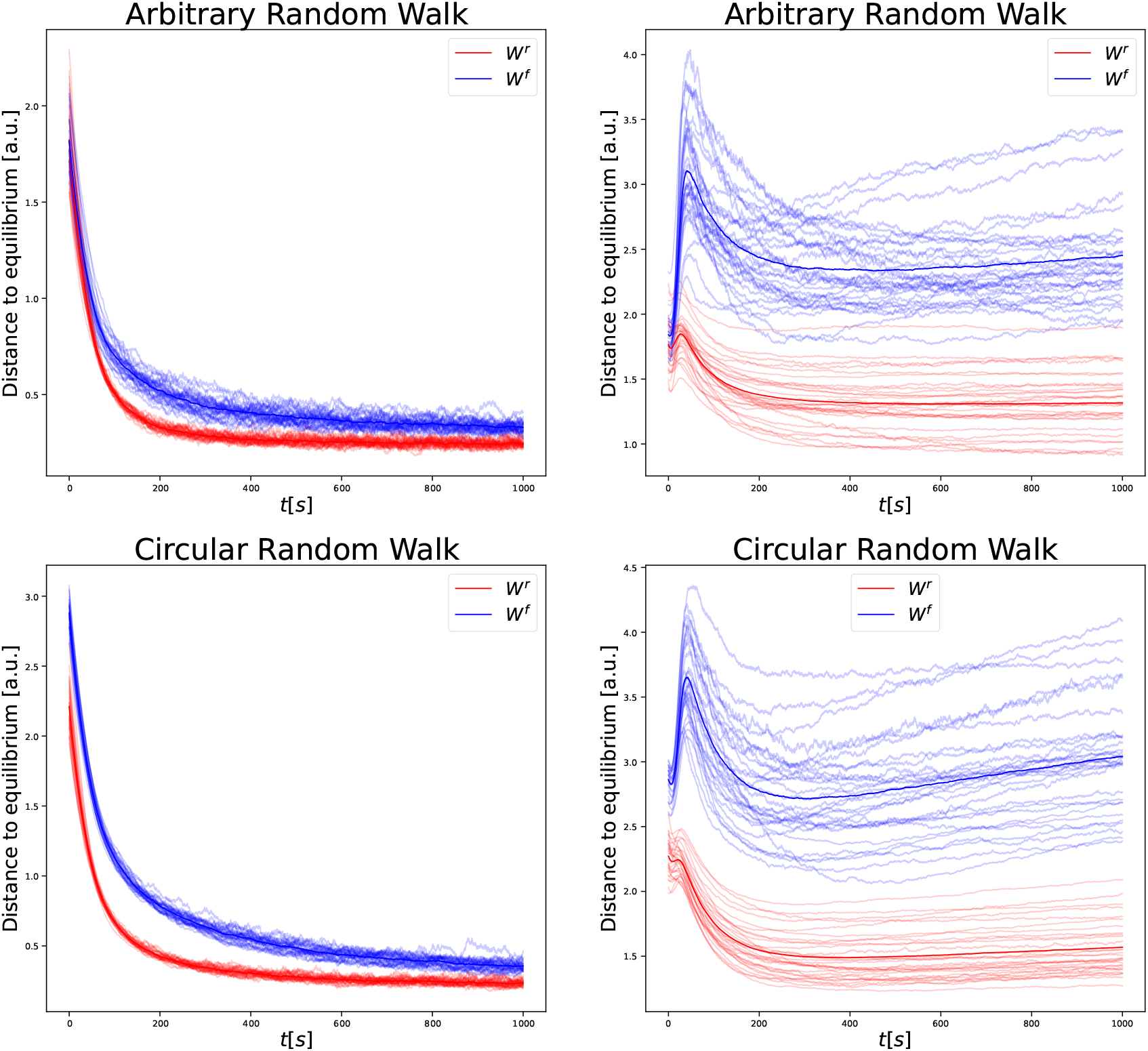
Convergence experiments with different activation functions. We conducted the same experiments as in Figure 3, with the activation functions (tanh,relu) from left to right.

**Figure H.2:**
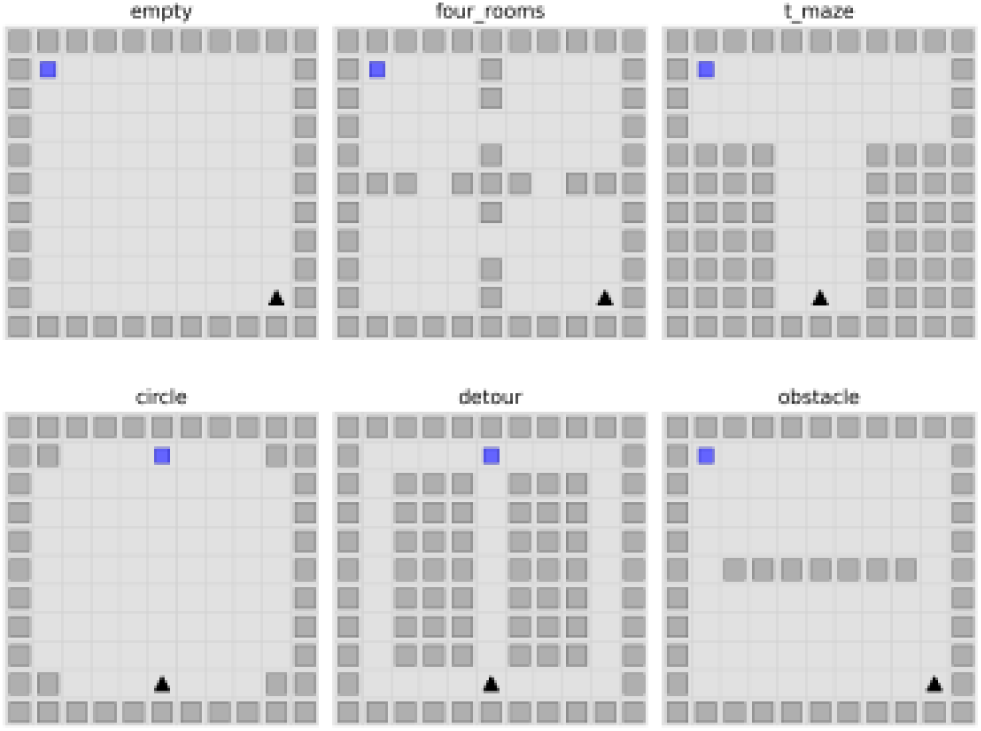
Grid world environments used in navigation tasks. Outline of the environments used to produce Figure 6. In all environments, agents started at random locations and could choose between four actions, corresponding to moving one step in the respective direction (or staying in place, when the supposed next position would be a wall). Plots were generated using Neuronav package Juliani et al. (2022).

**Figure H.3:**
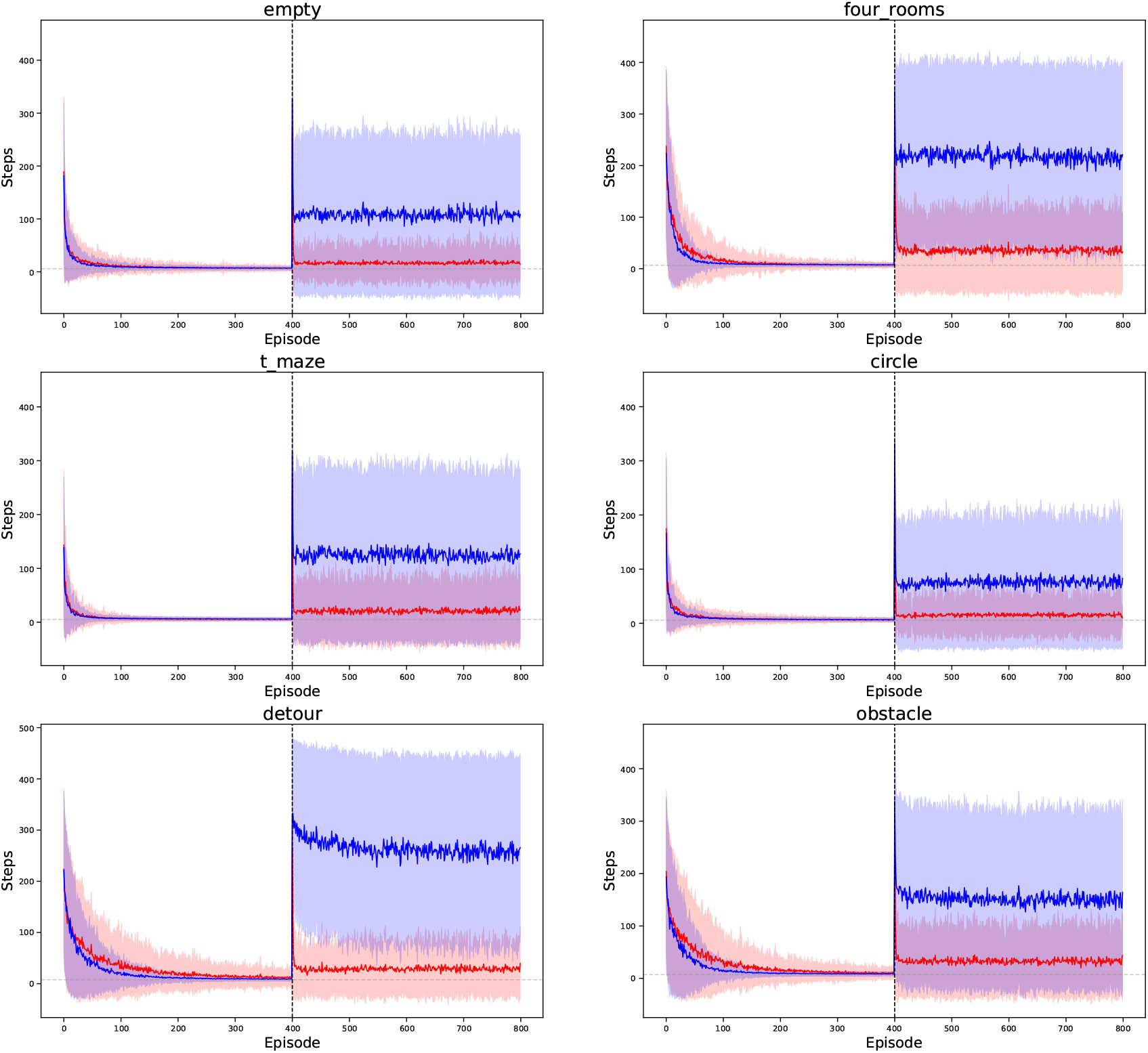
Generalization performance in individual environments with fixed number of episoded. Generalization performance in the individual environments which are averaged over to generate the left plot in Figure 6.

**Figure H.4:**
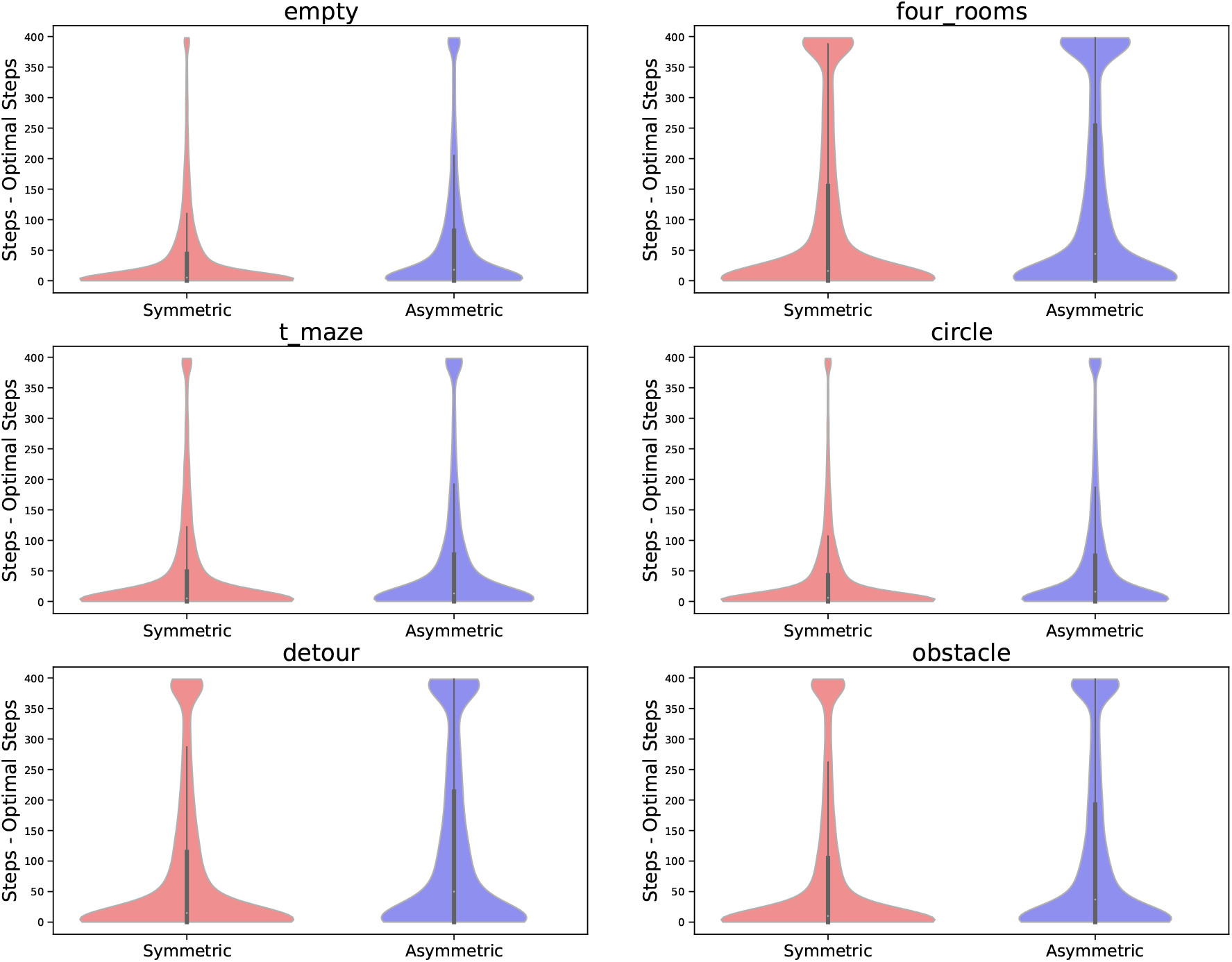
Generalization performance in individual environments with fixed accuracy criterion. Generalization performance in the individual environments which are averaged over to generate the right plot in Figure 6.

**Figure H.5:**
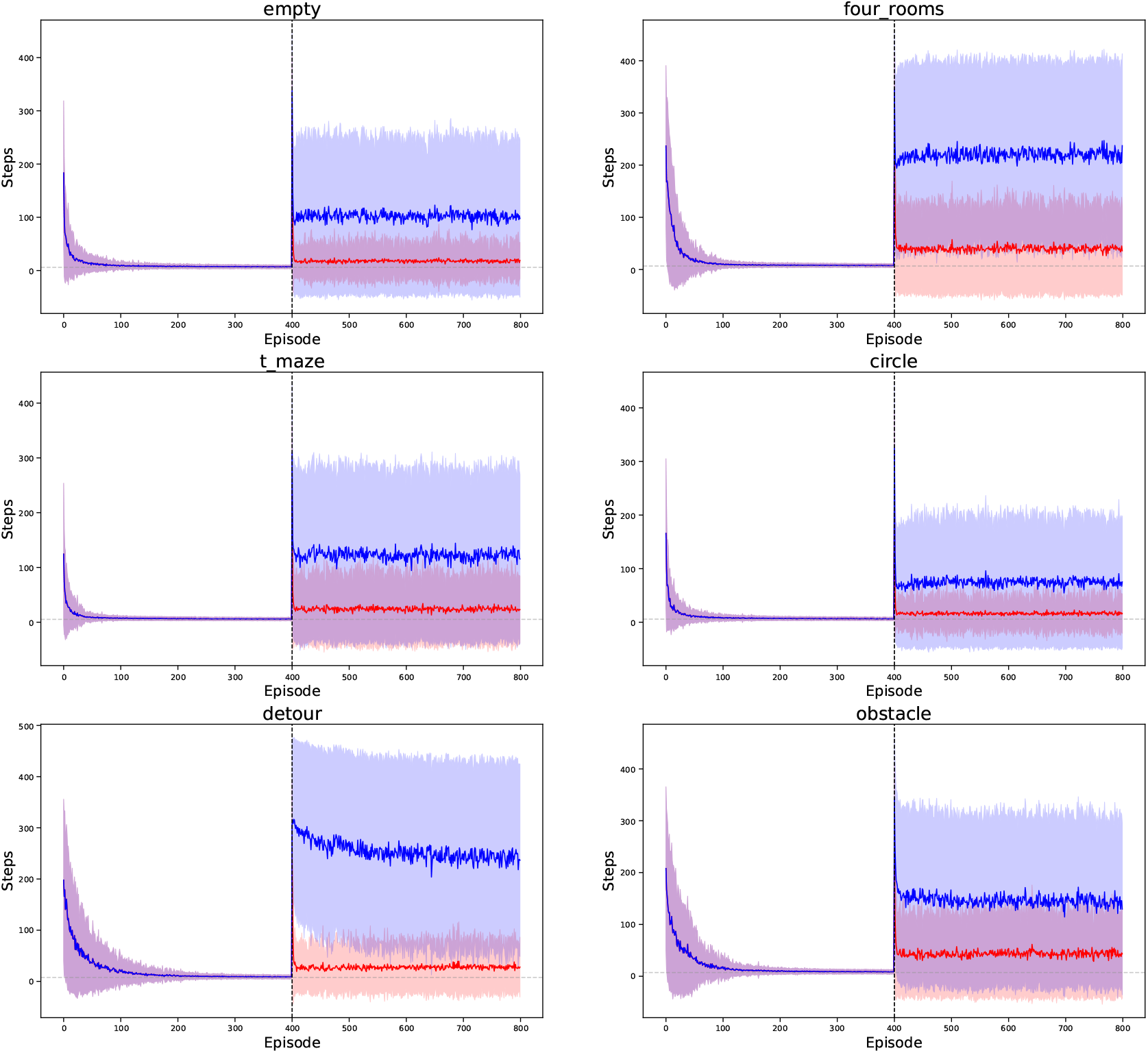
Generalization when seeing the same data while training. This figure is identical to Figure H.3, only that here we trained the symmetric agent on the transitions sampled by the asymmetric agent - hence both agents see exactly the same data before generalization.

**Figure H.6:**
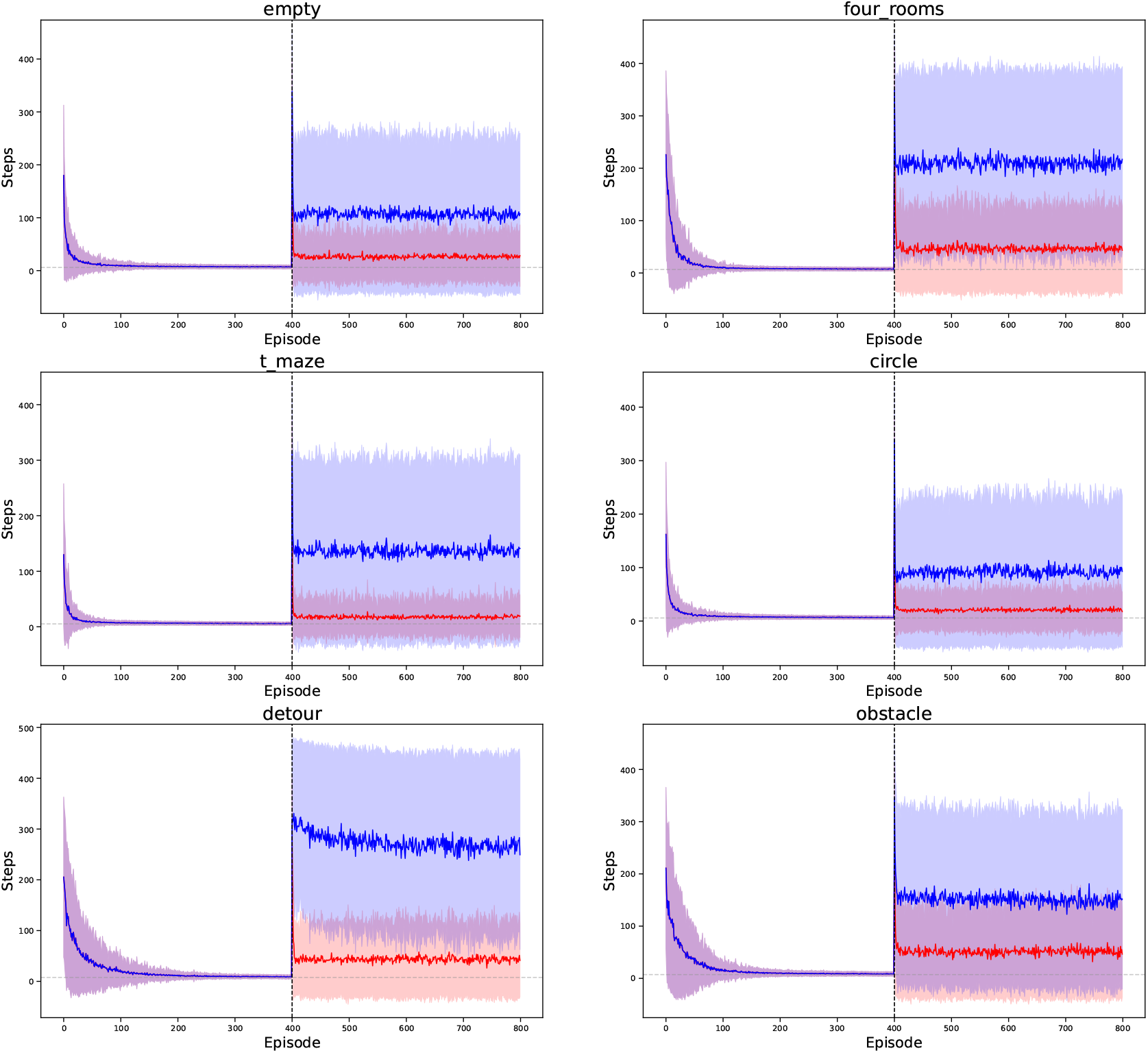
Generalization when seeing the same data while training and normalization. This figure is identical to Figure H.5, with the addition that now the updates to the SR are normalized, so that both agents take an equally sized update at each step..

**Figure H.7:**
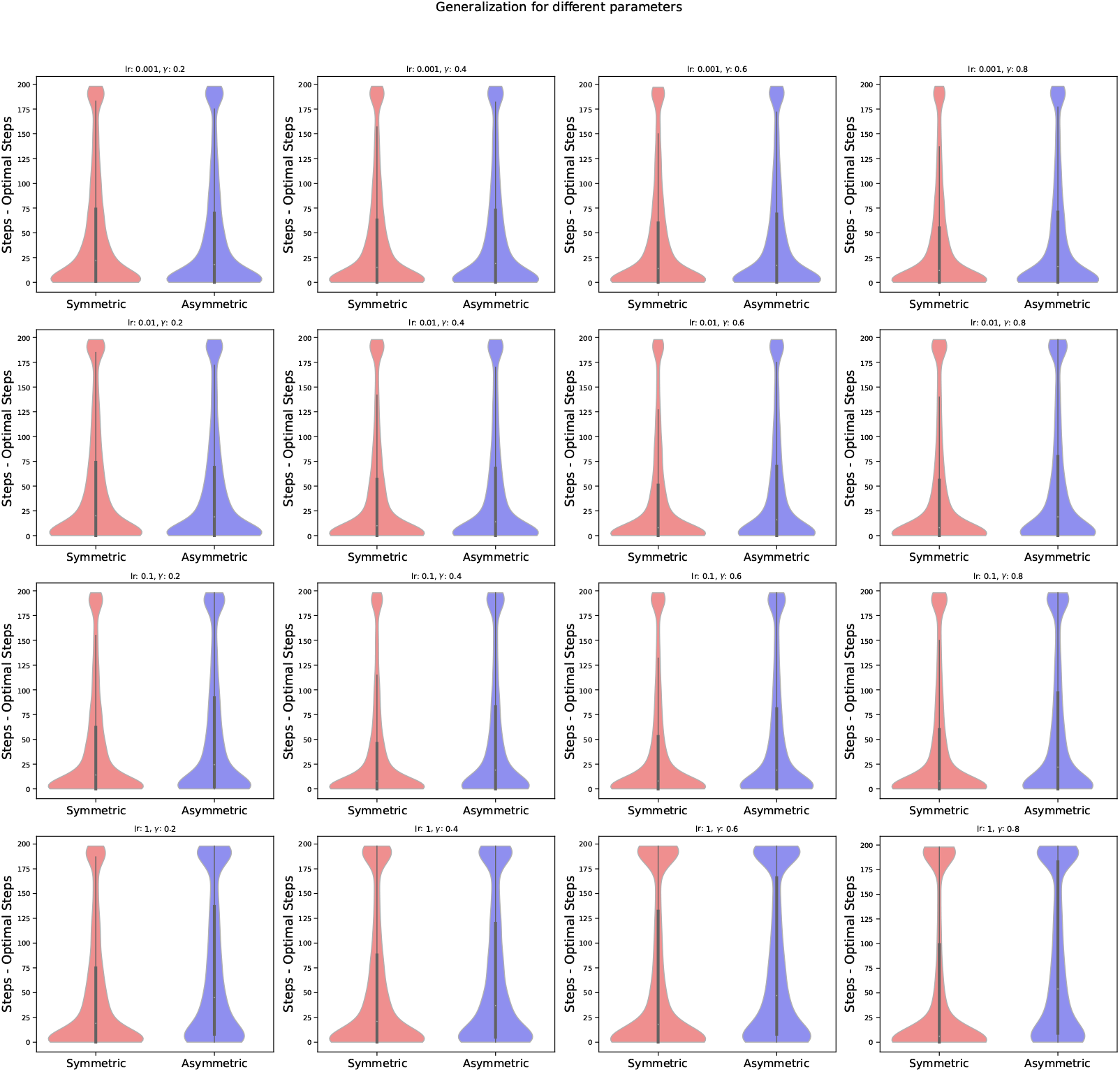
Generalization performance for varying choices of discount factors and learning rates. All experiments were conducted in the ‘empty’ environment.Note that in the case were the asymmetric agent outpasses the symmetric agent, generalization is considerably worse than in the regimes where it does not.

## References

Barreto, A., Borsa, D., Quan, J., Schaul, T., Silver, D., Hessel, M., Mankowitz, D., Zidek, A., and Munos, R. (2018). Transfer in deep reinforcement learning using successor features and generalised policy improvement. In International Conference on Machine Learning, pages 501–510. PMLR.

Barreto, A., Dabney, W., Munos, R., Hunt, J. J., Schaul, T., van Hasselt, H. P., and Silver, D. (2017). Successor features for transfer in reinforcement learning. Advances in neural information processing systems, 30.

Behrens, T. E., Muller, T. H., Whittington, J. C., Mark, S., Baram, A. B., Stachenfeld, K. L., and Kurth-Nelson, Z. (2018). What is a cognitive map? organizing knowledge for flexible behavior. Neuron, 100(2):490–509.

Bi, G.-q. and Poo, M.-m. (1998). Synaptic modifications in cultured hippocampal neurons: dependence on spike timing, synaptic strength, and postsynaptic cell type. Journal of neuroscience, 18(24):10464–10472.

Bono, J., Zannone, S., Pedrosa, V., and Clopath, C. (2023). Learning predictive cognitive maps with spiking neurons during behavior and replays. Elife, 12:e80671.

Bottini, R. and Doeller, C. F. (2020). Knowledge across reference frames: Cognitive maps and image spaces. Trends in Cognitive Sciences, 24(8):606–619.

Brzosko, Z., Mierau, S. B., and Paulsen, O. (2019). Neuromodulation of spiketiming-dependent plasticity: past, present, and future. Neuron, 103(4):563– 581.

Brzosko, Z., Schultz, W., and Paulsen, O. (2015). Retroactive modulation of spike timing-dependent plasticity by dopamine. elife, 4:e09685.

Canto, C. B., Wouterlood, F. G., Witter, M. P., et al. (2008). What does the anatomical organization of the entorhinal cortex tell us? Neural plasticity, 2008.

Carvalho, W., Tomov, M. S., de Cothi, W., Barry, C., and Gershman, S. J. (2024). Predictive representations: building blocks of intelligence. arXiv preprint 2402.06590.

Chung, F. (2005). Laplacians and the cheeger inequality for directed graphs. Annals of Combinatorics, 9:1–19.

Chung, F. R. (1996). Laplacians of graphs and cheeger’s inequalities. Combinatorics, Paul Erdos is Eighty, 2(157-172):13–2.

Constantinescu, A. O., O’Reilly, J. X., and Behrens, T. E. (2016). Organizing conceptual knowledge in humans with a gridlike code. Science, 352(6292):1464–1468.

Corneil, D. S. and Gerstner, W. (2015). Attractor network dynamics enable preplay and rapid path planning in maze–like environments. Advances in neural information processing systems, 28.

Dayan, P. (1993). Improving generalization for temporal difference learning: The successor representation. Neural computation, 5(4):613–624.

Dayan, P. and Sejnowski, T. J. (1994). Td (λ) converges with probability 1. Machine Learning, 14:295–301.

De Cothi, W. and Barry, C. (2020). Neurobiological successor features for spatial navigation. Hippocampus, 30(12):1347–1355.

De Cothi, W., Nyberg, N., Griesbauer, E.-M., Ghanamé, C., Zisch, F., Lefort, J. M., Fletcher, L., Newton, C., Renaudineau, S., Bendor, D., et al. (2022). Predictive maps in rats and humans for spatial navigation. Current Biology, 32(17):3676–3689.

Debanne, D., Gähwiler, B. H., and Thompson, S. M. (1999). Heterogeneity of synaptic plasticity at unitary CA3–CA1 and CA3–CA3 connections in rat hippocampal slice cultures. Journal of Neuroscience, 19(24):10664–10671.

Diba, K. and Buzsáki, G. (2007). Forward and reverse hippocampal place-cell sequences during ripples. Nature neuroscience, 10(10):1241–1242.

Dong, C., Madar, A. D., and Sheffield, M. E. (2021). Distinct place cell dynamics in CA1 and CA3 encode experience in new environments. Nature communications, 12(1):2977. This work is licensed under the Creative Commons Attribution 4.0 International License. To view a copy of this license, visit http://creativecommons.org/licenses/by/4.0/.

Dordek, Y., Soudry, D., Meir, R., and Derdikman, D. (2016). Extracting grid cell characteristics from place cell inputs using non-negative principal component analysis. Elife, 5:e10094.

Eichenbaum, H., Dudchenko, P., Wood, E., Shapiro, M., and Tanila, H. (1999). The hippocampus, memory, and place cells: is it spatial memory or a memory space? Neuron, 23(2):209–226.

Fang, C., Aronov, D., Abbott, L., and Mackevicius, E. L. (2023). Neural learning rules for generating flexible predictions and computing the successor representation. Elife, 12:e80680.

Fiete, I. R., Burak, Y., and Brookings, T. (2008). What grid cells convey about rat location. Journal of Neuroscience, 28(27):6858–6871.

Foster, D. J. and Wilson, M. A. (2006). Reverse replay of behavioural sequences in hippocampal place cells during the awake state. Nature, 440(7084):680–683.

Friston, K. (2002). Functional integration and inference in the brain. Progress in neurobiology, 68(2):113–143.

Garvert, M. M., Saanum, T., Schulz, E., Schuck, N. W., and Doeller, C. F. (2023). Hippocampal spatio-predictive cognitive maps adaptively guide reward generalization. Nature Neuroscience, 26(4):615–626.

Geerts, J. P., Chersi, F., Stachenfeld, K. L., and Burgess, N. (2020). A general model of hippocampal and dorsal striatal learning and decision making. Proceedings of the National Academy of Sciences, 117(49):31427–31437.

George, T., Stachenfeld, K., Barry, C., Clopath, C., and Fukai, T. (2023a). A generative model of the hippocampal formation trained with theta driven local learning rules. In Thirty-seventh Conference on Neural Information Processing Systems.

George, T. M., de Cothi, W., Clopath, C., Stachenfeld, K., and Barry, C. (2022). Ratinabox: A toolkit for modelling locomotion and neuronal activity in continuous environments. bioRxiv, pages 2022–08.

George, T. M., de Cothi, W., Stachenfeld, K. L., and Barry, C. (2023b). Rapid learning of predictive maps with stdp and theta phase precession. Elife, 12:e80663.

Gershman, S. J. (2018). The successor representation: its computational logic and neural substrates. Journal of Neuroscience, 38(33):7193–7200.

Givan, R., Dean, T., and Greig, M. (2003). Equivalence notions and model minimization in markov decision processes. Artificial Intelligence, 147(1-2):163– 223.

Hafting, T., Fyhn, M., Molden, S., Moser, M.-B., and Moser, E. I. (2005). Microstructure of a spatial map in the entorhinal cortex. Nature, 436(7052):801– 806.

Haga, T. and Fukai, T. (2018). Recurrent network model for learning goaldirected sequences through reverse replay. Elife, 7:e34171.

Hasselmo, M. E. (2006). The role of acetylcholine in learning and memory. Current opinion in neurobiology, 16(6):710–715.

Higgins, I., Racanière, S., and Rezende, D. (2022). Symmetry-based representations for artificial and biological general intelligence. Frontiers in Computational Neuroscience, 16:836498.

Huang, Y. and Rao, R. P. (2011). Predictive coding. Wiley Interdisciplinary Reviews: Cognitive Science, 2(5):580–593.

Ishizuka, N., Weber, J., and Amaral, D. G. (1990). Organization of intrahip-pocampal projections originating from ca3 pyramidal cells in the rat. Journal of comparative neurology, 295(4):580–623.

Johns, J. and Mahadevan, S. (2007). Constructing basis functions from directed graphs for value function approximation. In Proceedings of the 24th international conference on Machine learning, pages 385–392.

Jost, J. and Mulas, R. (2019). Cheeger-like inequalities for the largest eigenvalue of the graph laplace operator. arXiv preprint 1910.12233.

Juliani, A., Barnett, S., Davis, B., Sereno, M., and Momennejad, I. (2022). Neuro-nav: a library for neurally-plausible reinforcement learning. arXiv preprint 2206.03312.

Khona, M. and Fiete, I. R. (2022). Attractor and integrator networks in the brain. Nature Reviews Neuroscience, 23(12):744–766.

Knierim, J. J. (2015). The hippocampus. Current Biology, 25(23):R1116–R1121.

Kumaran, D. and McClelland, J. L. (2012). Generalization through the recurrent interaction of episodic memories: a model of the hippocampal system. Psychological review, 119(3):573.

Kushner, H. J. and Clark, D. S. (2012). Stochastic approximation methods for constrained and unconstrained systems, volume 26. Springer Science & Business Media.

Lee, I., Rao, G., and Knierim, J. J. (2004a). A double dissociation between hippocampal subfields: differential time course of CA3 and CA1 place cells for processing changed environments. Neuron, 42(5):803–815.

Lee, I., Yoganarasimha, D., Rao, G., and Knierim, J. J. (2004b). Comparison of population coherence of place cells in hippocampal subfields CA1 and CA3. Nature, 430(6998):456–459.

Leutgeb, S., Leutgeb, J. K., Treves, A., Moser, M.-B., and Moser, E. I. (2004). Distinct ensemble codes in hippocampal areas CA3 and CA1. Science, 305(5688):1295–1298.

Levin, D. A. and Peres, Y. (2017). Markov chains and mixing times, volume 107. American Mathematical Soc.

Machado, M. C., Bellemare, M. G., and Bowling, M. (2017a). A laplacian framework for option discovery in reinforcement learning. In International Conference on Machine Learning, pages 2295–2304. PMLR.

Machado, M. C., Rosenbaum, C., Guo, X., Liu, M., Tesauro, G., and Campbell, M. (2017b). Eigenoption discovery through the deep successor representation. arXiv preprint 1710.11089.

Mahadevan, S. and Maggioni, M. (2007). Proto-value functions: A laplacian framework for learning representation and control in markov decision processes. Journal of Machine Learning Research, 8(10).

Markram, H., Gerstner, W., and Sjöström, P. J. (2011). A history of spiketiming-dependent plasticity. Frontiers in synaptic neuroscience, 3:4.

Markram, H., Lübke, J., Frotscher, M., and Sakmann, B. (1997). Regulation of synaptic efficacy by coincidence of postsynaptic aps and epsps. Science, 275(5297):213–215.

Mavor-Parker, A. N., Sargent, M. J., Banino, A., Griffin, L. D., and Barry, C. (2022). A simple approach for state-action abstraction using a learned mdp homomorphism. arXiv preprint 2209.06356.

Mehta, M. R., Barnes, C. A., and McNaughton, B. L. (1997). Experience-dependent, asymmetric expansion of hippocampal place fields. Proceedings of the National Academy of Sciences, 94(16):8918–8921.

Mercatali, G., Freitas, A., and Garg, V. (2022). Symmetry-induced disentanglement on graphs. Advances in Neural Information Processing Systems, 35:31497–31511.

Mishra, R. K., Kim, S., Guzman, S. J., and Jonas, P. (2016). Symmetric spike timing-dependent plasticity at CA3–CA3 synapses optimizes storage and recall in autoassociative networks. Nature communications, 7(1):11552.

Mizuseki, K., Royer, S., Diba, K., and Buzsáki, G. (2012). Activity dynamics and behavioral correlates of ca3 and ca1 hippocampal pyramidal neurons. Hippocampus, 22(8):1659–1680.

Momennejad, I., Russek, E. M., Cheong, J. H., Botvinick, M. M., Daw, N. D., and Gershman, S. J. (2017). The successor representation in human reinforcement learning. Nature human behaviour, 1(9):680–692.

Namboodiri, V. M. K. and Stuber, G. D. (2021). The learning of prospective and retrospective cognitive maps within neural circuits. Neuron, 109(22):3552– 3575.

Nitsch, A., Garvert, M. M., Bellmund, J. L., Schuck, N. W., and Doeller, C. F. (2023). Grid-like entorhinal representation of an abstract value space during prospective decision making. bioRxiv, pages 2023–08.

Norris, J. R. (1998). Markov chains. Number 2. Cambridge university press.

O’Keefe, J. and Nadel, L. (1978). The hippocampus as a cognitive map. Hippocampus, 3:570.

Ollivier, Y. (2018). Approximate temporal difference learning is a gradient descent for reversible policies. arXiv preprint 1805.00869.

Pawlak, V., Wickens, J. R., Kirkwood, A., and Kerr, J. N. (2010). Timing is not everything: neuromodulation opens the stdp gate. Frontiers in synaptic neuroscience, 2:146.

Penny, W. D., Zeidman, P., and Burgess, N. (2013). Forward and backward inference in spatial cognition. PLoS computational biology, 9(12):e1003383.

Piray, P. and Daw, N. D. (2021). Linear reinforcement learning in planning, grid fields, and cognitive control. Nature communications, 12(1):4942.

Rao, R. P. and Ballard, D. H. (1999). Predictive coding in the visual cortex: a functional interpretation of some extra-classical receptive-field effects. Nature neuroscience, 2(1):79–87.

Roth, E. D., Yu, X., Rao, G., and Knierim, J. J. (2012). Functional differences in the backward shifts of CA1 and CA3 place fields in novel and familiar environments. PloS one, 7(4):e36035.

Russek, E. M., Momennejad, I., Botvinick, M. M., Gershman, S. J., and Daw, N. D. (2017). Predictive representations can link model-based reinforcement learning to model-free mechanisms. PLoS computational biology, 13(9):e1005768.

Samsonovich, A. and McNaughton, B. L. (1997). Path integration and cognitive mapping in a continuous attractor neural network model. Journal of Neuroscience, 17(15):5900–5920.

Scoville, W. B. and Milner, B. (1957). Loss of recent memory after bilateral hippocampal lesions. Journal of neurology, neurosurgery, and psychiatry, 20(1):11.

Seabrook, E. and Wiskott, L. (2023). A tutorial on the spectral theory of markov chains. Neural Computation, 35(11):1713–1796.

Seol, G. H., Ziburkus, J., Huang, S., Song, L., Kim, I. T., Takamiya, K., Huganir, R. L., Lee, H.-K., and Kirkwood, A. (2007). Neuromodulators control the polarity of spike-timing-dependent synaptic plasticity. Neuron, 55(6):919–929.

Sprekeler, H. (2011). On the relation of slow feature analysis and laplacian eigenmaps. Neural computation, 23(12):3287–3302.

Squire, L. R., Stark, C. E., and Clark, R. E. (2004). The medial temporal lobe. Annu. Rev. Neurosci., 27:279–306.

Stachenfeld, K. L., Botvinick, M., and Gershman, S. J. (2014). Design principles of the hippocampal cognitive map. Advances in neural information processing systems, 27.

Stachenfeld, K. L., Botvinick, M. M., and Gershman, S. J. (2017). The hippocampus as a predictive map. Nature neuroscience, 20(11):1643–1653.

Sugisaki, E., Fukushima, Y., Tsukada, M., and Aihara, T. (2011). Cholinergic modulation on spike timing-dependent plasticity in hippocampal ca1 network. Neuroscience, 192:91–101.

Sutton, R. S. and Barto, A. G. (2018). Reinforcement learning: An introduction. MIT press.

Theves, S., Neville, D. A., Fernández, G., and Doeller, C. F. (2021). Learning and representation of hierarchical concepts in hippocampus and prefrontal cortex. Journal of Neuroscience, 41(36):7675–7686.

Van der Pol, E., Kipf, T., Oliehoek, F. A., and Welling, M. (2020a). Plannable approximations to mdp homomorphisms: Equivariance under actions. arXiv preprint 2002.11963.

Van der Pol, E., Worrall, D., van Hoof, H., Oliehoek, F., and Welling, M. (2020b). Mdp homomorphic networks: Group symmetries in reinforcement learning. Advances in Neural Information Processing Systems, 33:4199–4210.

Vértes, E. and Sahani, M. (2019). A neurally plausible model learns successor representations in partially observable environments. Advances in Neural Information Processing Systems, 32.

Von Luxburg, U. (2007). A tutorial on spectral clustering. Statistics and computing, 17:395–416.

Whittington, J. C., Muller, T. H., Mark, S., Chen, G., Barry, C., Burgess, N., and Behrens, T. E. (2020). The tolman-eichenbaum machine: unifying space and relational memory through generalization in the hippocampal formation. Cell, 183(5):1249–1263.

Wittenberg, G. M. and Wang, S. S.-H. (2006). Malleability of spike-timing-dependent plasticity at the ca3–ca1 synapse. Journal of Neuroscience, 26(24):6610–6617.

Witter, M. P. (2007). Intrinsic and extrinsic wiring of ca3: indications for connectional heterogeneity. Learning & memory, 14(11):705–713.

Wu, Y., Tucker, G., and Nachum, O. (2018). The laplacian in rl: Learning representations with efficient approximations. arXiv preprint 1810.04586.

Yu, C., Burgess, N., Sahani, M., and Gershman, S. (2023). Successor-predecessor intrinsic exploration. arXiv preprint 2305.15277.

Yu, C., Li, D., Hao, J., Wang, J., and Burgess, N. (2021). Learning state representations via retracing in reinforcement learning. arXiv preprint 2111.12600.

Zannone, S., Brzosko, Z., Paulsen, O., and Clopath, C. (2018). Acetylcholine-modulated plasticity in reward-driven navigation: a computational study. Scientific reports, 8(1):9486.

Zhang, K. (1996). Representation of spatial orientation by the intrinsic dynamics of the head-direction cell ensemble: a theory. Journal of Neuroscience, 16(6):2112–2126.

Zhang, T., Rosenberg, M., Jing, Z., Perona, P., and Meister, M. (2021). Endotaxis: A neuromorphic algorithm for mapping, goal-learning, navigation, and patrolling. bioRxiv, pages 2021–09.

